# Mitochondrial pyruvate transport regulates presynaptic metabolism and neurotransmission

**DOI:** 10.1101/2024.03.20.586011

**Authors:** Anupama Tiwari, Jongyun Myeong, Arsalan Hashemiaghdam, Hao Zhang, Xianfeng Niu, Marissa A. Laramie, Marion I. Stunault, Jasmin Sponagel, Gary Patti, Leah Shriver, Vitaly Klyachko, Ghazaleh Ashrafi

## Abstract

Glucose has long been considered the primary fuel source for the brain. However, glucose levels fluctuate in the brain during sleep, intense circuit activity, or dietary restrictions, posing significant metabolic stress. Here, we demonstrate that the mammalian brain utilizes pyruvate as a fuel source, and pyruvate can support neuronal viability in the absence of glucose. Nerve terminals are sites of metabolic vulnerability within a neuron and we show that mitochondrial pyruvate uptake is a critical step in oxidative ATP production in hippocampal terminals. We find that the mitochondrial pyruvate carrier is post-translationally modified by lysine acetylation which in turn modulates mitochondrial pyruvate uptake. Importantly, our data reveal that the mitochondrial pyruvate carrier regulates distinct steps in synaptic transmission, namely, the spatiotemporal pattern of synaptic vesicle release and the efficiency of vesicle retrieval, functions that have profound implications for synaptic plasticity. In summary, we identify pyruvate as a potent neuronal fuel and mitochondrial pyruvate uptake as a critical node for the metabolic control of synaptic transmission in hippocampal terminals.

**HIGHLIGHTS:**

- Serum pyruvate is taken up by the brain and efficiently oxidized in the TCA cycle.
- The mitochondrial pyruvate carrier (MPC) is essential for presynaptic energy metabolism.
- Acetylation of the MPC complex modulates mitochondrial pyruvate uptake.
- MPC activity regulates the release and retrieval of synaptic vesicles in nerve terminals.

## INTRODUCTION

The brain requires a constant, ready source of energy to function properly. When metabolic processes supporting energy generation fail, for example secondary to ischemic stroke, or uncontrolled diabetes, loss of cognitive function follows rapidly^1,2^. Paradoxically, even in a healthy brain, access to glucose, its canonical energy source, is relatively unreliable. In fact, glucose concentration in the brain interstitial fluid is estimated to be low (∼1mM)^3^, which is further depleted by bouts of high neuronal activity^4,5^, during fasting, and sleep^6^. Neurons consume ∼40% of total ATP in the cortex to sustain synaptic transmission^7^, the signaling process underpinning cognition. The fluctuating levels of glucose in the brain, coupled with limited glycogen storage in this organ, suggests that neurons would have to rely on alternative energy sources to sustain synaptic transmission^8^. These alternative fuels primarily include amino acids, ketone bodies, pyruvate and its derivative lactate.

There are two major routes for neurons to acquire pyruvate: the blood supply, and local lactate production by neighboring astrocytes. Although the physiological relevance of pyruvate delivery to the brain remains unclear, it is known that endothelial cells and astrocytic end feet forming the blood-brain barrier express monocarboxylate transporters which can transport pyruvate from the blood to brain interstitial fluid^9^. Indeed, magnetic resonance imaging demonstrated that pyruvate can enter the human brain from the circulation^10^, supporting the notion that pyruvate could serve as an energy source for the brain^11–14^. Yet, the extent to which pyruvate can support neuronal functions, and the molecular mechanisms of pyruvate oxidation in neurons remain poorly understood.

Pyruvate is an oxidative fuel that is broken down via the tricarboxylic (TCA) cycle in the mitochondrial matrix. Once taken up into neuronal cytosol by monocarboxylate transporters, pyruvate must traverse mitochondrial inner and outer membranes to gain access to TCA enzymes. The voltage-dependent anion channel (VDAC) makes mitochondrial outer membrane largely permeable to metabolites, such as pyruvate^15^. However, pyruvate passage through the inner mitochondrial membrane is facilitated by the mitochondrial pyruvate carrier (MPC), a multimeric complex of two subunits: MPC1 and MPC2^16,17^. By regulating pyruvate entry to the mitochondrial matrix and its access to the TCA cycle, MPC mediates a crucial branch point in energy metabolism. The physiological importance of MPC is further highlighted by its implication in several human pathologies, including cancer, cardiac hypertrophy, diabetes, and microcephaly^18^. Furthermore, there is also growing interest in targeting MPC for treatment of neurological disorders such as Parkinson’s disease^19,20^. However, few studies have examined MPC function in the nervous system^21,22^, particularly in the metabolic control of synaptic transmission.

Mathematical models suggest that mitochondrial pyruvate uptake is a limiting step in stimulation of oxidative ATP synthesis^23^. Indeed, yeast cells upregulate pyruvate oxidation under aerobic growth conditions through alternative expression of MPC isoforms with enhanced transport kinetics^24^. However, this mechanism is not conserved in mammalian cells, and it is not known how MPC activity is modulated in energetically demanding cells like neurons. In the present study, we investigated pyruvate metabolism in the brain and discovered its critical role in the regulation of synaptic transmission. We demonstrated that mitochondrial pyruvate transport is essential for ATP production in nerve terminals and regulates distinct steps in the synaptic vesicle cycle. We further uncovered a post-translational mechanism for regulation of mitochondrial pyruvate transport. Altogether, our study establishes that mitochondrial pyruvate uptake in nerve terminals is precisely modulated to ensure the metabolic plasticity of neurotransmission.

## RESULTS

### Pyruvate is efficiently oxidized in intact brain and is a metabolic fuel for neuronal cultures

It has been previously shown that acute supplementation of cultured neurons or brain slices with pyruvate can sustain the energetics of neurotransmission^12–14^. However, it is not clear to what extent chronic pyruvate supply can support neuronal survival in the absence of glucose, and whether pyruvate can serve as a bona fide fuel source for the intact brain. To examine pyruvate metabolism in the rodent brain, we performed intravenous infusion of ^13^C_3_-labeled pyruvate into the jogular vein of C57BL/6 mice that were briefly fasted (∼5 hours), and analyzed the metabolic fate of pyruvate in serum, cortex and cerebellum, as well as liver using liquid chromatography and mass spectrometry (LC-MS) (**Fig. 1A**). We detected labeled pyruvate and lactate in the brain (cortex and cerebellum combined) suggesting that these metabolites cross the blood brain barrier to enter the interstitial fluid (**Fig. 1C**). We then investigated the metabolic fate of pyruvate in various tissues by quantifying ^13^C-labeled intermediates of pyruvate metabolism in gluconeogenesis and oxidative phosphorylation (i.e. the TCA cycle) (**Fig. 1B**). Consistent with the well-established role of the liver in gluconeogenesis, ^13^C labeling of glucose and gluconeogenic intermediates were higher in the liver than the brain (**Fig 1C and Table S1**). In contrast, labeled TCA intermediates were significantly enriched in the brain compared to serum and the liver indicating preferential oxidation of pyruvate in the brain (**Fig 1C and Table S1**). Interestingly, ^13^C-labeling of TCA intermediates in the brain was much more pronounced than either pyruvate or lactate, suggesting that oxidative breakdown of pyruvate is highly favorable in the brain. Altogether, our data demonstrates that pyruvate enters the brain from the circulation where it undergoes oxidative phosphorylation via the TCA cycle. These findings indicate that pyruvate is a bona fide fuel source for the brain under physiological conditions.

**Figure 1.**
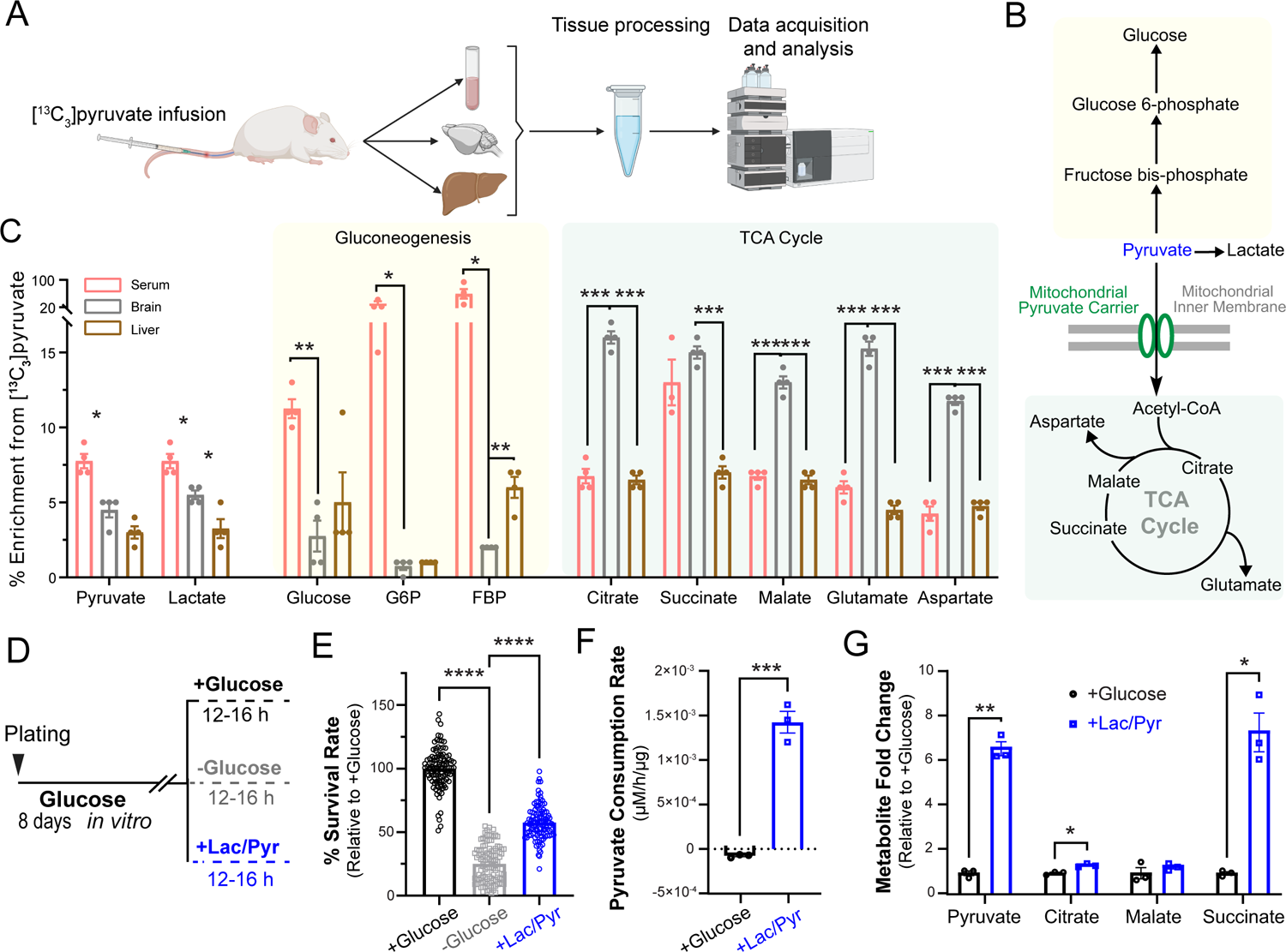
Pyruvate is efficiently oxidized in intact brain and is a metabolic fuel for primary neuronal cultures. (A) Schematic for metabolic tracing of intravenously perfused ^13^C_3_ pyruvate in mouse serum, brain, and liver using LC-MS. Created with Biorender.com. (B) Pathways of pyruvate metabolism including gluconeogenesis and oxidation via the tricarboxylic acid (TCA) cycle. (C) Enrichment of ^13^C-labeled intermediates of gluconeogenesis and the TCA cycle in different tissues. n = 4 mice. (D) Schematic for survival analysis and metabolic profiling of primary neurons at DIV8 supplied with various fuel types, with glucose (5mM), no glucose, or an equimolar mix of lactate and pyruvate (5mM each) denoted as lac/pyr. (E) Survival rate of neurons relative to control (+glucose). n = 108 (wells), 3 (cultures). (F) Pyruvate consumption rate (μM/hr/μg of lysate) in neurons treated as in (E). n = 3 (wells). (G) Steady-state levels of TCA metabolites in neurons treated as in (D) plotted as fold change relative to +glucose condition = 3 (wells). Unpaired t-test (C, F, G), and One-way ANOVA (E).*p < 0.05; ** p < 0.01, ***p < 0.001, ****p < 0.0001.

Our *in vivo* metabolomic analysis captures the combined metabolic fates of pyruvate across all cell types comprising brain tissue. To examine pyruvate metabolism specifically in a less heterogeneous cellular population, we cultured primary rat cortical neurons and investigated whether pyruvate can sustain neuronal viability in the absence of glucose. In this paradigm, neurons cultured under standard conditions were divided into three groups: supplied with glucose (5mM), supplied with an equimolar mixture of lactate and pyruvate (5mM each) (denoted as lac/pyr) in the absence of glucose, or provided with neither glucose nor lac/pyr (i.e. fuel deprivation) for 12-16 hours before assessing viability (**Fig. 1D**). While fuel deprivation dramatically reduced neuronal survival to ∼20%, supplementation with lac/pyr in the absence of glucose could maintain neuronal survival at ∼50% of the glucose (control) condition (**Fig. 1E and Table S2**). This finding demonstrates that pyruvate supply can partially sustain neuronal viability even in the absence of glucose. To investigate neuronal pyruvate metabolism in more detail, primary cortical neurons supplied with glucose or pyruvate were analyzed with LC–MS-based targeted metabolomics. While neurons supplied with glucose released pyruvate into the culture media, neurons supplied with pyruvate were observed to taken up extracellular pyruvate (**Fig. 1F**), supporting its potential use as an energy source. Measurement of steady state levels of intracellular pyruvate and other TCA cycle intermediates with LC-MS revealed upregulation of their pool size in neurons supplied with pyruvate, consistent with higher pyruvate flux through the TCA cycle (**Fig. 1G and Table S2**). These findings suggest that neurons efficiently oxidize pyruvate as an alternative energy source to glucose and neuronal pyruvate metabolism can partially sustain neuronal viability.

### Mitochondrial pyruvate uptake is essential for energy metabolism in nerve terminals

The MPC complex is expressed at high levels in the nervous system^21^, however little is known about its subcellular distribution in neurons. To determine whether MPC is present in nerve terminals where there is substantial energy demand for synaptic transmission, dissociated hippocampal neurons were immunostained with antibodies against MPC subunits (MPC1 and MPC2) and the presynaptic protein - vesicular glutamate transporter 1(vGLUT1). Both subunits were present in neuronal cell bodies (**Fig. 2A and 2C**), but they also colocalized with vGLUT1, indicating the expression of MPC1 and MPC2 in presynaptic terminals (**Fig. 2B and 2D**).

**Figure 2.**
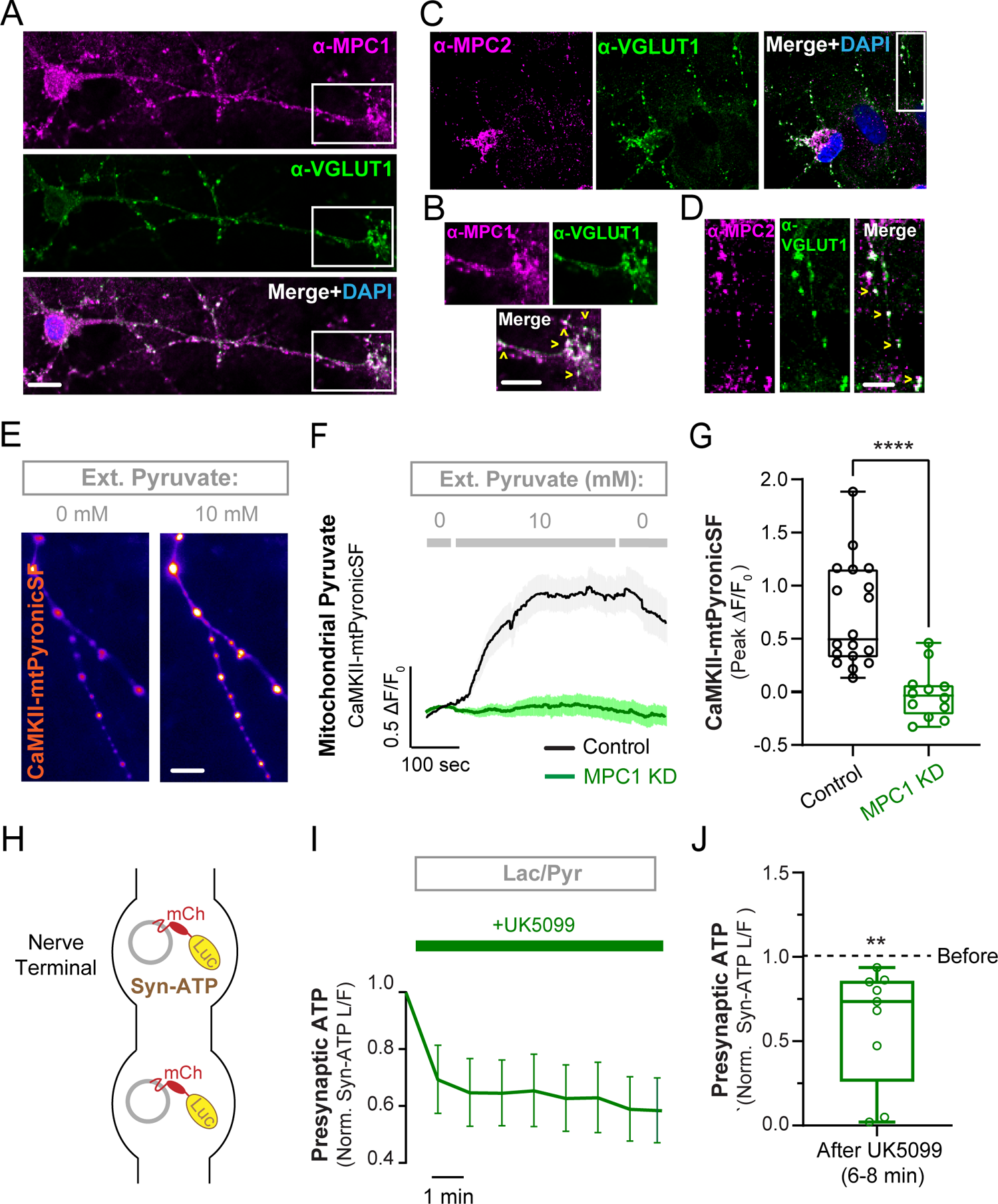
Mitochondrial pyruvate carrier is essential for pyruvate metabolism in nerve terminals. **(A, C)** Co-immunostaining of hippocampal neurons with antibodies against MPC1 (A) or MPC2 (C) and the presynaptic protein vGLUT1. Scale bars, 10 µm. **(B, D)** Magnification of the boxed areas in panels A and C. Scale bars, 10 µm (B), 5 µm (D). (E) Representative images of axonal mitochondria expressing a mitochondrial pyruvate sensor, CamKII-mito-PyronicSF, showing an increase in fluorescence intensity with perfusion of 10mM extracellular pyruvate, compared to 0mM. Scale bar, 10 µm. (F) Average traces of CamKII-mito-PyronicSF(ΔF/F_0_) showing inhibition of mitochondrial pyruvate uptake in MPC1 KD neurons. n =12 –18(neurons). (G) Maximal CamKII-mito-PyronicSF fluorescence response in 10mM pyruvate as determined from traces in panel F. (H) Schematic of the genetically encoded ATP indicator Syn-ATP expressed in nerve terminals. (I) Presynaptic ATP traces in terminals supplied with lactate and pyruvate (lac/pyr) after incubation with the MPC inhibitor UK5099, normalized to pre-incubation levels. n =10 (neurons). (J) Presynaptic ATP level after application of MPC inhibitor UK5099 normalized to ATP level before. The box-whisker plots denote median (line), 25^th^-75^th^ percentile (box), and min-max (whiskers). Mann-Whitney U test (G), One sample t-test (I).

To determine whether the MPC complex mediates mitochondrial pyruvate uptake in presynaptic mitochondria, we developed a quantitative optical assay for mitochondrial pyruvate uptake using the fluorescent pyruvate sensor PyronicSF. In this sensor, the bacterial transcription factor PdhR is fused with cyclically permuted GFP (cpGFP) and targeted to the mitochondrial matrix with a COXVIII mitochondrial targeting sequence^23^. To improve expression of PyronicSF in neurons, a construct using the CaMKII promoter^25^ was developed, denoted as CaMKII-mtPyronicSF. In neuronal axons expressing the sensor, perfusion of 10mM extracellular pyruvate led to a significant increase in mitochondrial fluorescence (**Fig. 2E-2G**), consistent with the import of pyruvate into the mitochondrial matrix. The extracellular pyruvate concentration of 10mM was selected based on the affinity and dynamic range of the parental PyronicSF sensor^23^. Depletion of neuronal MPC1 with shRNA (denoted as MPC1 KD) completely abolished CaMKII-mtPyronicSF fluorescence response to extracellular pyruvate, thus validating proper subcellular localization and specificity of the reporter (**Fig. 2F and 2G**). This finding also indicated that the MPC complex is essential for pyruvate accumulation in axonal mitochondria. The efficiency of MPC1 depletion was confirmed by qPCR of *mpc1* mRNA in cortical neuronal cultures (**Fig. S1A**). Since neuronal MPC2 protein was previously found to be unstable when MPC1 was depleted^22^, we concluded that MPC1 KD effectively ablates both components of the complex.

The presynaptic abundance of MPC subunits led us to investigate the effects of MPC inhibition on presynaptic ATP levels in neurons supplied with an equimolar mixture of lactate and pyruvate and no glucose. In terminals expressing the genetically encoded ATP indicator Syn-ATP^26^ (**Fig. 2H**), pharmacological inhibition of MPC with the compound UK5099^27^ caused significant depletion of presynaptic ATP within a few minutes of application, indicating the critical role of MPC in pyruvate oxidation (**Fig. 2I and 2J**). Taken together these results demonstrate that the MPC complex is present in presynaptic mitochondria and is essential for oxidative pyruvate metabolism in nerve terminals.

### Lysine acetylation of MPC complex modulates mitochondrial pyruvate uptake

Lysine acetylation is a reversible post-translational modification that plays a major role in modulation of mitochondrial function^28^. While it is thought that mitochondrial proteins are acetylated non-enzymatically, the removal of acetyl groups is catalyzed by mitochondrial sirtuins, primarily Sirtuin 3 (Sirt3), an NAD^+^-dependent deacetylase. We previously discovered that neuronal glucose deprivation induces Sirt3 expression, which in turn stimulates oxidative metabolism in nerve terminals^29^. Interestingly, MPC2 has been found to be hyper-acetylated in the liver of *Sirt3^-/-^* mice^30^ and in diabetic heart^31^, implicating acetylation as a potential mechanism for modulation of MPC activity.

These previous findings prompted us to ask whether acetylation of mitochondrial proteins by Sirt3 regulates the pyruvate transport activity of the MPC complex. To examine mitochondrial pyruvate uptake in a higher throughput manner than in neuronal axons, the optical pyruvate sensor mtPyronicSF was expressed in HEK293 cells and extracellular pyruvate concentration was changed from 0mM to 10mM, triggering a reversible rise in the fluorescence intensity of mtPyronicSF consistent with mitochondrial pyruvate accumulation (**Fig. 3A and 3B**). Similar to neuronal axons (**Fig. 2E-G**), this response was fully blocked by shRNA-mediated depletion of MPC1 (MPC1 KD) (**Fig. 3A and 3B**) which was confirmed by qPCR of *mpc1* mRNA (**Fig. S2A**). Importantly, shRNA-mediated Sirt3 KD significantly blunted mitochondrial pyruvate accumulation, implicating Sirt3 in modulation of mitochondrial pyruvate transport. The efficiency of Sirt3 KD was confirmed by western blotting for Sirt3 protein (**Fig. S2B and S2C**). Since the proton gradient across the inner mitochondrial membrane provides the driving force for pyruvate transport by the MPC complex^32^, we confirmed that MPC1 KD or Sirt3 KD did not alter mitochondrial membrane potential in HEK293 cells by staining with the mitochondrial membrane dye TMRM (**Fig. S2D**).

**Figure 3.**
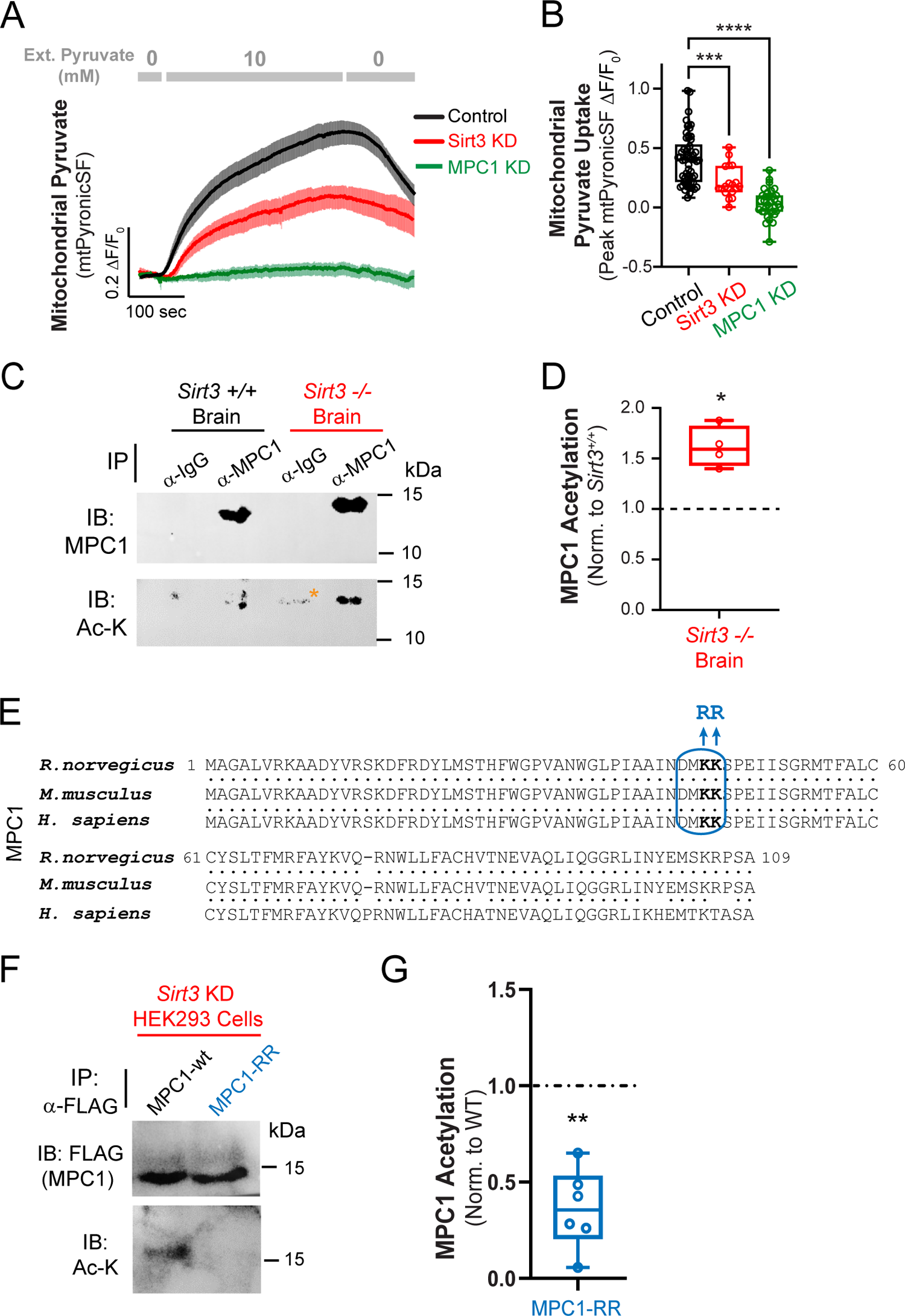
MPC acetylation regulates mitochondrial pyruvate uptake and synaptic transmission. (A) Average traces of mito-PyronicSF expressed in HEK293 cells and treated with varying extracellular pyruvate concentrations, showing differential mitochondrial pyruvate accumulation in control HEK293 cells, and cells expressing shRNA against Sirt3 (Sirt3 KD) or MPC1 (MPC1 KD). (B) Peak values mito-PyronicSF (ΔF/F_0_) in response to 10mM pyruvate, as determined from traces in panel A. n = 18 – 58 fields of view (FOV), 5-6 cells/FOV. (C) Immunoprecipitation (IP) of endogenous MPC1 from *Sirt3^+/+^*and *Sirt3^-/-^*mouse brain lysates with antibodies against MPC1 and rabbit IgG (control) followed by immunoblotted (IB) with antibodies against MPC1 (top) and Ac-K (bottom). Asterix denotes a non-specific band in the IgG IP. (D) Intensity of Ac-K band normalized to MPC1 band from panel C, expressed relative to *Sirt3^+/+^*control. n = 4 (IP blots). Raw intensities of Ac-K normalized to MPC1 in *Sirt3^+/+^*and *Sirt3^-/-^*brains were used for statistical comparison. **(E)** Alignment of rat (*R. norvegicus*), mouse (*M. musculus*), and human (*H. sapiens*) MPC1 protein sequences showing conservation of an acetylation motif (boxed in blue). A non-acetylatable form was constructed by mutation of K45/46 to R45/R46. (E) Immunoprecipitation of wildtype MPC1 (MPC1-wt) or acetyl mutant (MPC1-RR) from Sirt3-deficient (KD) HEK293 cells, immunoblotted for FLAG (top) and Ac-K (bottom). (F) Intensity of Ac-K band normalized to FLAG intensity from panel E, expressed relative to wt-MPC1.n = 6 (IP blots). Kruskal-Wallis test (B), Paired t-test (D), One sample t-test (G).

Our data suggest that Sirt3 regulates pyruvate entry into the mitochondrial matrix, potentially through post-translational modification of MPC subunits. To further examine this possibility, we investigated whether Sirt3 abundance impacts the acetylation state of the MPC complex in the brain. We focused on MPC1, because unlike MPC2, acetylation of this subunit has not been reported in proteomics studies and remains poorly explored^31^. MPC1 was immunoprecipitated from *Sirt3^+/+^*(*Sirt3^fl/fl^*) and *Sirt3^-/-^* mouse^33^ brain tissues and the immunoprecipitated (IP) fraction was immunoblotted with a pan-acetyl-lysine antibody revealing hyperacetylation of MPC1 in *Sirt3^-/-^* brain as compared to *Sirt3^+/+^* tissue (**Fig.3C and 3D**). The specificity of MPC1 IP band was confirmed with an antibody against mouse IgG. Our results thus demonstrate that MPC1 is a substrate for Sirt3-mediated deacetylation in the brain.

To further confirm MPC1 acetylation, we sought to map the acetylated lysine residues. Bioinformatic analysis of lysine acetylation in mitochondrial proteins from rat brain and brown fat has identified a consensus sequence motif for lysine acetylation (D-X-X-AcK)^34^ which is conserved in MPC1 sequences from rats, mice, and humans (**Fig. 4E**, boxed in blue). To confirm if the conserved lysines in MPC1 were indeed acetylated as predicted, we constructed a double mutant of FLAG-tagged MPC1 in which K45 and K46 motif sites were replaced with arginine (denoted as MPC1-RR), thus preserving electrostatic charge while blocking acetylation. To facilitate detection of MPC1 acetylation, FLAG-tagged (wildtype) MPC1-wt or MPC1-RR were expressed and immunoprecipitated from Sirt3-deficient HEK293 cells which display enhanced acetylation of mitochondrial proteins^35,36^. Sirt3 knockdown with CRISPR-Cas9 gene editing was confirmed with immunoblotting HEK293 lysates from successive passages with an antibody against Sirt3 (**Fig. S2E**). While both MPC1-wt and MPC1-RR were pulled down with anti-FLAG antibody, immunoblotting the IP fraction with a pan-acetyl-lysine antibody revealed significant reduction of MPC1-RR acetylation as compared to (wildtype) MPC1-wt (**Fig. 4F and 4G**). This result confirms K45 and K46 as the primary acetylated residues in MPC1. Altogether, our findings suggest that Sirt3 regulates MPC1 acetylation in HEK293 cells and brain tissue, and that it also modulates mitochondrial pyruvate uptake.

**Figure 4.**
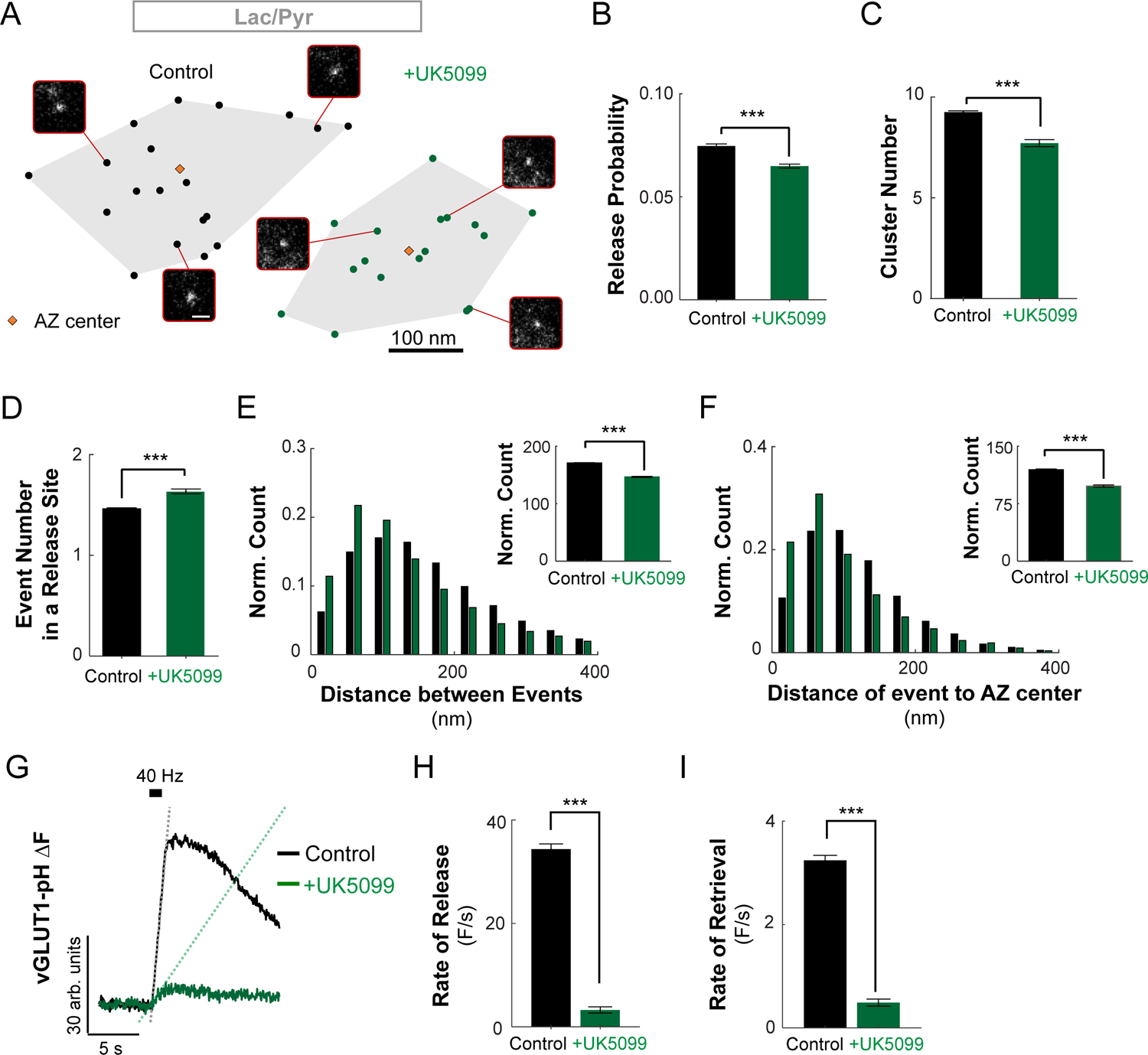
Mitochondrial pyruvate uptake regulates spatiotemporal properties of SV release and retrieval. **(A)** Two examples of AZs with the localization of release events without (left) or with (right) UK5099. Representative images of individual events evoked at 1Hz are shown for each AZ. **(B)** Average release probability of events evoked by 1Hz stimulation for 200 seconds in the two conditions. Sample number (cultures/coverslips/synapses): Control: 9/36/1163, UK5099: 3/9/23 **(C, D)** Average number of clusters/release sites in an individual AZ (C) and number of events detected per release site (D) in the two conditions. Sample number: control: 9/36/1254/11925, UK5099: 3/9/220/1705 for cultures/coverslips/synapses/release sites. (E) Distribution of distances between consecutive events over time. The inset shows the average distance from the distribution below. Sample number: control: 3/9/15084, UK5099: 3/9/2786 for cultures/dishes/events. (F) Distribution of distances from release events to the center of the AZ. The inset shows the average distance from the distributions below. Sample number: control: 3/9/15084, UK5099: 3/9/2786 for cultures/dishes/events. (G) Sample traces of vesicle release evoked by high frequency stimulation of 50 AP at 40 Hz in the two conditions. **(H,I)** Rates of vesicle exocytosis **(H)** and endocytosis **(I)** in the two conditions calculated by linear fitting of the rise and decay components of the train responses during high-frequency stimulation. Sample number: control: 3/9/1163, UK5099: 3/9/230 for cultures/dishes/synapses. Two Sample t-test (B, C, D, E, F, H, and I).

### Mitochondrial pyruvate uptake regulates distinct steps in the synaptic vesicle cycle

The release and retrieval of SVs, collectively known as the SV cycle, is a critical process in neurotransmission and is highly susceptible to energetic perturbations^26^. Since our results suggest that mitochondrial pyruvate uptake is essential for maintaining presynaptic ATP levels (**Fig. 2**), and MPC activity is post-translationally modulated (**Fig. 3**), we investigated the how MPC function might impact specific steps in the SV cycle. To examine SV release at a single-vesicle level in individual hippocamal synapses in culture, we used our previously established near-TIRF imaging approach combined with well-established computational detection and localization methods^37–39^. We visualized vesicle release with vGLUT1-pHluorin^40^ which contains a pH-sensitive indicator pHluorin targeted to the synaptic vesicle lumen via fusion with vGLUT1.

Neurons were acutely supplied with an equimolar mixture of lactate and pyruvate (denoted as lac/pyr) replacing glucose to enhance metabolic reliance on mitochondrial OXPHOS as we previously reported^12^. Individual release events were evoked by single action potential (AP) stimulation at 1Hz for 200 seconds and then automatically detected and localized within the synaptic active zone (AZ) with ∼27 nm precision using computational detection and localization approaches, as previously described^37,38,41^. The functional AZ area was defined as a convoluted hull encompassing all events identified within an individual synapse, with the centroid defining the AZ center (**Fig. 4A**). This definition of the AZ area is consistent with AZ dimensions observed in ultrastructural studies by electron microscopy^41,42^. The detected SV release events were clustered to define the location of individual release sites within each AZ using a hierarchical clustering algorithm with a clustering diameter of 50 nm^41–43^.

Using this approach, we first determined if MPC inhibition affected the basal vesicle release probability (Pr). Acute pharmacological inhibition of MPC with UK5099 significantly reduced the Pr measured at 1 Hz stimulation (**Fig. 4B**). SV release is not randomly distributed across the AZ, but is characterized by a repeated utilization of several specialized release sites within the AZ^37,38,42–45^. Given the reduction in P_r_ and the energetic dependencies of SV release, we next asked whether the spatial organization of SV release is affected by MPC inhibition. We found that the number of vesicle release sites, denoted as cluster number, was significantly reduced (**Fig. 4C**), while repeated utilization of the same release sites increased with MPC inhibition (**Fig. 4D**). Furthermore, SV release events occurred at significantly shorter distances from each other (**Fig. 4E**), and the distance from the events to the AZ center was also significantly reduced when MPC was inhibited (**Fig. 4F**). These results suggest that mitochondrial pyruvate uptake during single AP firing regulates the spatial dynamics of SV release shifting release towards a subset of more centrally located sites, as depicted schematically (**Fig. 4A**).

The energetic demands of synaptic transmission scale with the duration and intensity of stimulation^46^. Therefore, we examined the impact of MPC inhibition on SV release and retrieval evoked by high-frequency stimulus trains of 50 AP at 40 Hz in hippocampal terminals supplied with lactate and pyruvate (**Fig. 4G**). We found that SV release was significantly impaired when MPC was acutely inhibited with UK5099 (**Fig.4H**). The retrieval of SVs after release is also known to be highly sensitive to energetic perturbations^12,14,26^. Indeed, we found that MPC inhibition dramatically slowed SV retrieval following stimulation by a train of 50 AP at 40 Hz (**Fig. 4I**). We confirmed that SV retrieval kinetics was similarly reduced following a longer stimulation paradigm (100 AP at 10 Hz) with application of UK5099 (**Fig. S3A and S3B**), or Zaprinast, another pharmacological inhibitor of MPC^47^ (**Fig. S3C and S3D**). Consistent with pharmacological inhibition, shRNA-mediated MPC1 KD significantly slowed SV retrieval following stimulation with 100 AP at 10 Hz, and this defect was fully rescued with expression of an shRNA-resistant MPC1 construct (**Fig. S3E and S3F**). Altogether, our data demonstrate that mitochondrial pyruvate transport by the MPC complex regulates the spatiotemporal properties of SV release and retrieval in nerve terminals, both during single AP firing and high-frequency trains.

## DISCUSSION

Glucose has long been considered the canonical energy source for the brain. Yet, glucose availability undergoes spatial and temporal fluctuations in active brain regions, and during sleep or fasting. Thus, the mammalian brain may have evolved to scavenge for alternative fuels such as pyruvate to protect against loss of cognitive function and ensure organismal survival in the face of unpredictable food supply. Here, using *in vivo* metabolomics and isotope tracing, we demonstrated that circulating blood delivers pyruvate to the brain where it is efficiently broken down by oxidative phosphorylation. We found that neurons can utilize pyruvate in the absence of glucose, and pyruvate metabolism in nerve terminals relies on mitochondrial pyruvate uptake by the MPC complex. In addition, mitochondrial pyruvate uptake is modulated by lysine acetylation of MPC subunits and MPC activity regulates the spatiotemporal properties of evoked SV release and retrieval in nerve terminals.

The remarkable capacity of the brain for pyruvate oxidation compared to other tissues may be explained by its steep energetic demands, as well as the catastrophic consequences of brain energetic failure. Neurons in the brain not only take up pyruvate from the circulation, but are also locally supplied with lactate derived from the glycolytic breakdown of glucose in neighboring astrocytes in a process known as the astrocyte-neuron lactate shuttle (ANLS) model^48^. The existence of multiple routes for delivery of pyruvate or lactate to neurons underscores the importance of this fuel type for neuronal metabolism. However, further work is needed to delineate the physiological roles of local versus systemic delivery of pyruvate to the brain, and their respective contributions to supporting neuronal function during dietary conditions or intense circuit activity.

A significant fraction of brain energy is directed to nerve terminals to support synaptic transmission. Here, we identify MPC-dependent pyruvate entry into the TCA cycle as a critical node in the metabolic regulation of the SV cycle. Indeed, we showed that mitochondrial pyruvate transport is crucial for the spatiotemporal control of SV release and subsequent SV retrieval in the absence of glucose. However, the question remains whether MPC function is also important for synaptic transmission in glucose-rich conditions. Our recent study demonstrated that pyruvate is critical SV release and retrieval in terminals supplied with glucose^39^. These findings support the notion that mitochondrial pyruvate uptake regulates synaptic transmission under a broad range of nutrient conditions.

Here, we show that the mitochondrial deacetylase Sirt3 modulates MPC1 deacetylation. The energetic needs of nerve terminals are highly variable, and regulation of pyruvate entry to the mitochondrial matrix via post-translational acetylation of MPC may serve as a molecular rheostat matching oxidative ATP synthesis with presynaptic energy demand. We previously showed that Sirt3 deacetylates a number of mitochondrial proteins during neuronal glucose deprivation. Therefore, we hypothesize that changes in Sirt3 expression in response to physiological stimuli, such as, glucose deprivation^29^, prolonged synaptic activity^49^, or in pathological conditions of diabetes^50^ and neurodegeneration^51^, may ultimately regulate neuronal pyruvate metabolism through MPC acetylation. Considering the critical role of MPC in modulation of synaptic transmission, it will be important for future studies to determine the effects of MPC acetylation on synaptic plasticity and cognitive performance under these conditions.

In the present study, we mapped two lysine acetylation sites (K45 and K46) in MPC1 and identified the functional significance of these modifications for mitochondrial pyruvate uptake. Although atomic structures of MPC subunits and the assembled complex are not available, structural predictions by AlphaFold localized K45 and K46 to a flexible loop connecting two transmembrane helices. However, further studies are needed to determine how lysine acetylation impacts MPC1 structural conformation to regulate pyruvate transport by this complex. Although we did not directly examine MPC2 in this study, hyperacetylation of MPC2 has been observed in diabetic mouse heart^31^. Like MPC1 mutants, acetyl mimetic mutations of MPC2 were shown to dampen oxidative metabolism in cardiac muscle, presumably by reducing mitochondrial pyruvate uptake^31^. Given that MPC1 and MPC2 are essential components of the same of transport complex in neurons and in other cell types, we anticipate that both subunits are similarly regulated by acetylation. Beyond lysine acetylation, there is also growing evidence for the reprogramming of oxidative metabolism by other forms of post-translational modifications of mitochondrial proteins, such as, phosphorylation^52^, O-GlcNAcylation^53^, and succinylation^54^. Therefore, it will be important to examine the regulation of neuronal MPC by these modifications and determine their physiological role in the metabolic plasticity of synaptic transmission.

Pyruvate has been shown to be critical for memory formation^55^, particularly when local glucose levels decline in active circuits during memory tasks^5^. Furthermore, the activity of pyruvate dehydrogenase, which catalyzes the first step in pyruvate oxidation, strongly correlates with the intensity of neuronal firing^56^. However, an important unresolved question is to what extent pyruvate metabolism varies across different brain regions, such as, the hippocampus, cerebellum, or cortex, and whether these regions differ in their metabolic plasticity in usage of fuel types, such as, glucose, pyruvate, or ketone bodies. Understanding the metabolic plasticity of brain function in molecular detail has profound clinical implications for human physiology. Indeed, metabolic reprograming through MPC inhibition has emerged as an insulin sensitizing strategy for treatment of type II diabetes and has been shown to attenuate neurodegeneration in some Parkinson’s disease models^19,20^. Given our finding that mitochondrial pyruvate uptake regulates synaptic transmission, it is critical to elucidate the short-term or long-term effects of MPC-modulating drugs on cognitive function in healthy individuals and patients suffering from metabolic and neurodegenerative diseases.

## ACKNOWLEDGEMENTS

We thank B. Finck and K. McCommis for help with MPC1 immunoprecipitation, C. Bergom for Sirt3^+/+^ and Sirt3^-/-^ mice, and P. Verma for CRISPR/Ca9 gene editing. We also thank Washington University Center for Cellular Imaging and the Genome Technology Access Center at the McDonnell Genome Institute, which is partially supported by NCI Cancer Center Support Grant #P30 CA91842 to the Siteman Cancer Center and by ICTS/CTSA Grant# UL1TR002345 from the National Center for Research Resources (NCRR), and NIH Roadmap for Medical Research. This work was funded by the Washington University Institute of Clinical and Translational Sciences (G.A.) which is, in part, supported by the NIH/National Center for Advancing Translational Sciences (NCATS), the Klingenstein-Simons Fellowship in Neuroscience (G.A.), the Whitehall Foundation (G.A.), NIGMS R35GM147222 (G.A.), and NINDS R35 NS111596 (V.K.). The authors declare no competing financial interests.

## AUTHOR CONTRIBUTIONS

Anupama Tiwari: Investigation, formal analysis, visualization, and writing (original draft). Jongyun Myeong: Investigation, formal analysis, and visualization. Arsalan Hashemiaghdam: Investigation, and formal analysis. Hao Zhang: Investigation, and formal analysis. Xianfeng Niu: Investigation, and formal analysis. Marion I. Stunault: Investigation, formal analysis, and visualization. Jasmin Sponagel: formal analysis, and visualization. Gary Patti: Conceptualization, supervision, and funding acquisition. Leah Shriver: Conceptualization, supervision, and funding acquisition. Vitaly Klyachko: Conceptualization, funding acquisition, supervision, and writing (original draft, review, and editing). Ghazaleh Ashrafi: Conceptualization, funding acquisition, supervision, and writing (original draft, review, and editing).

## KEY RESOURCES TABLE

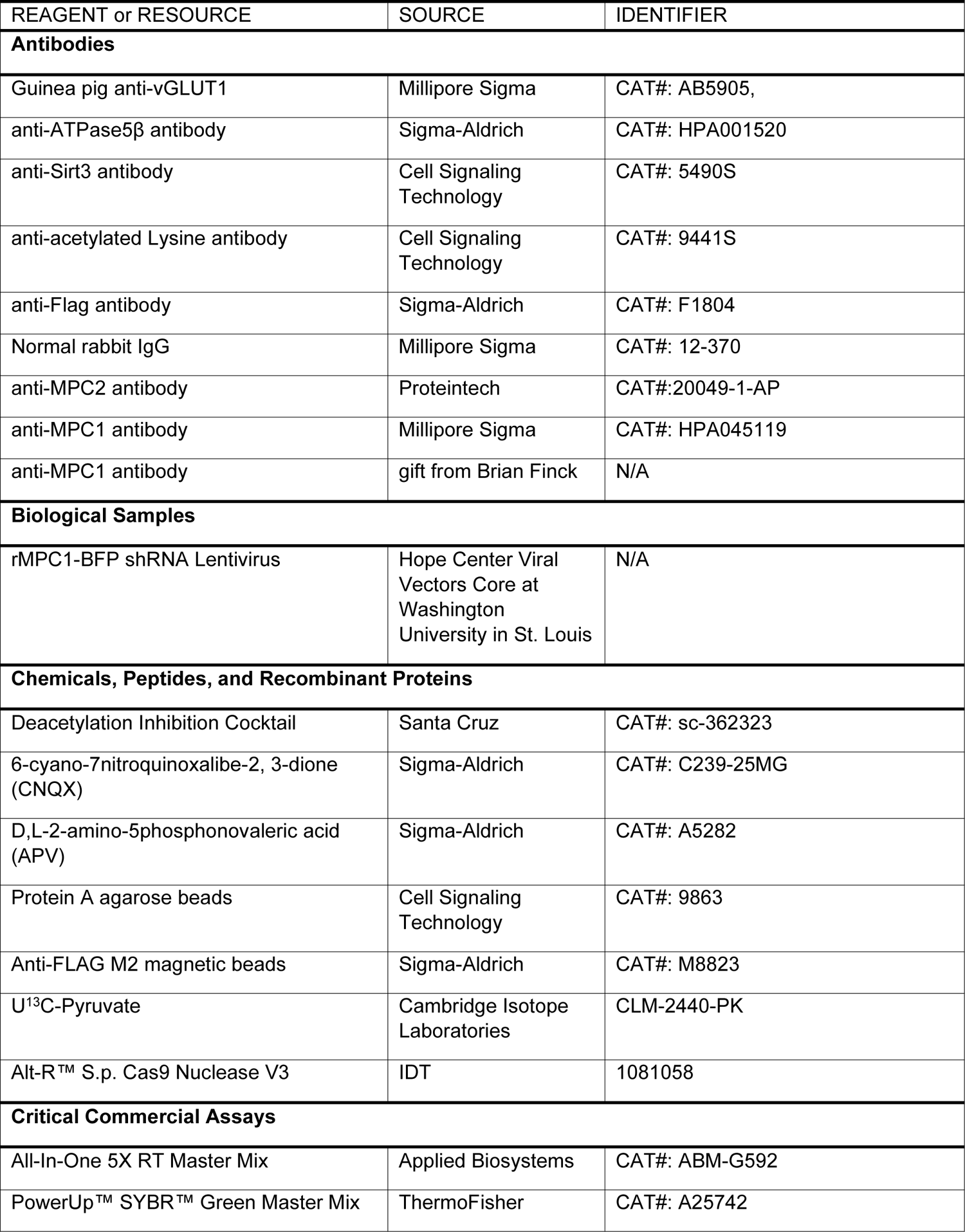
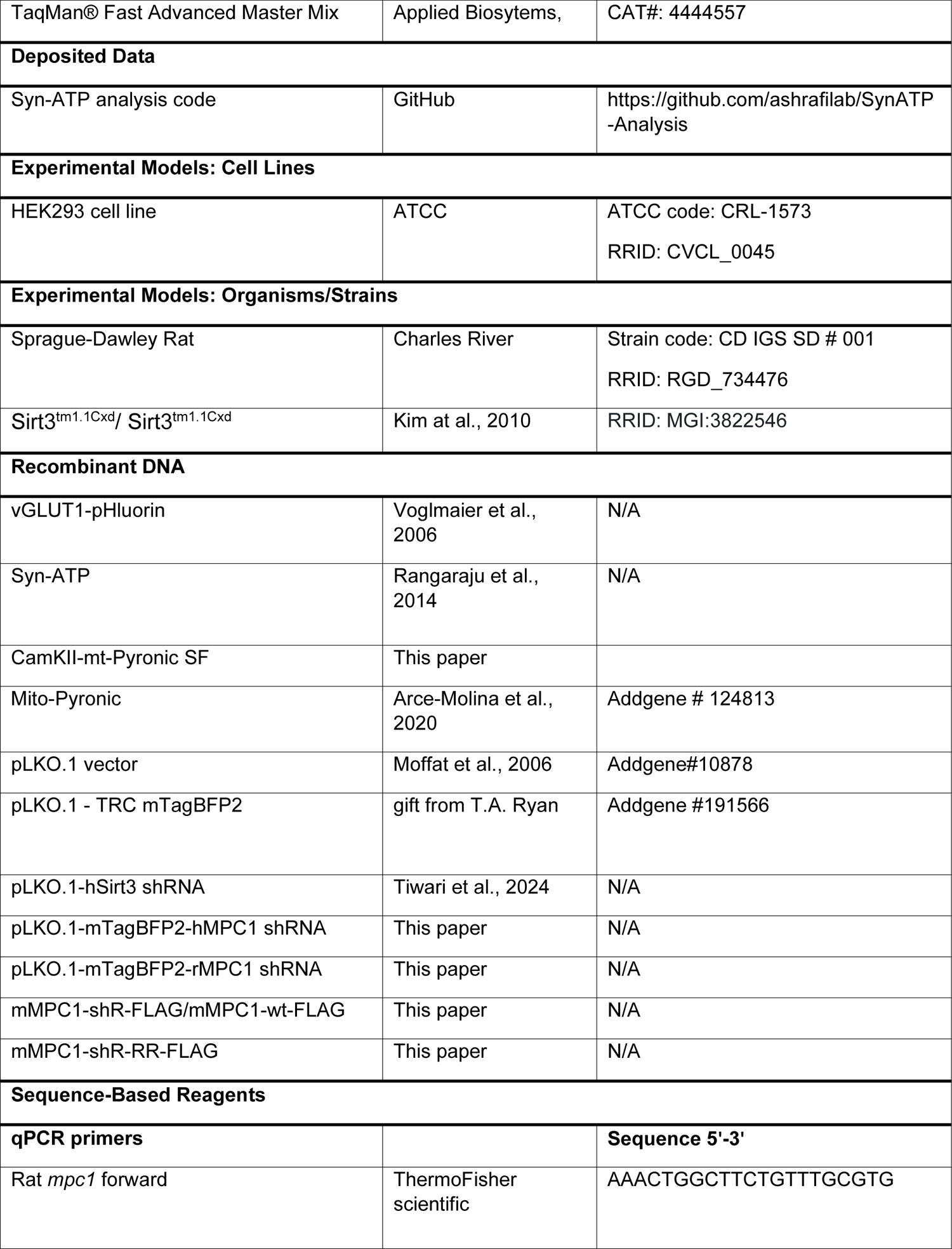
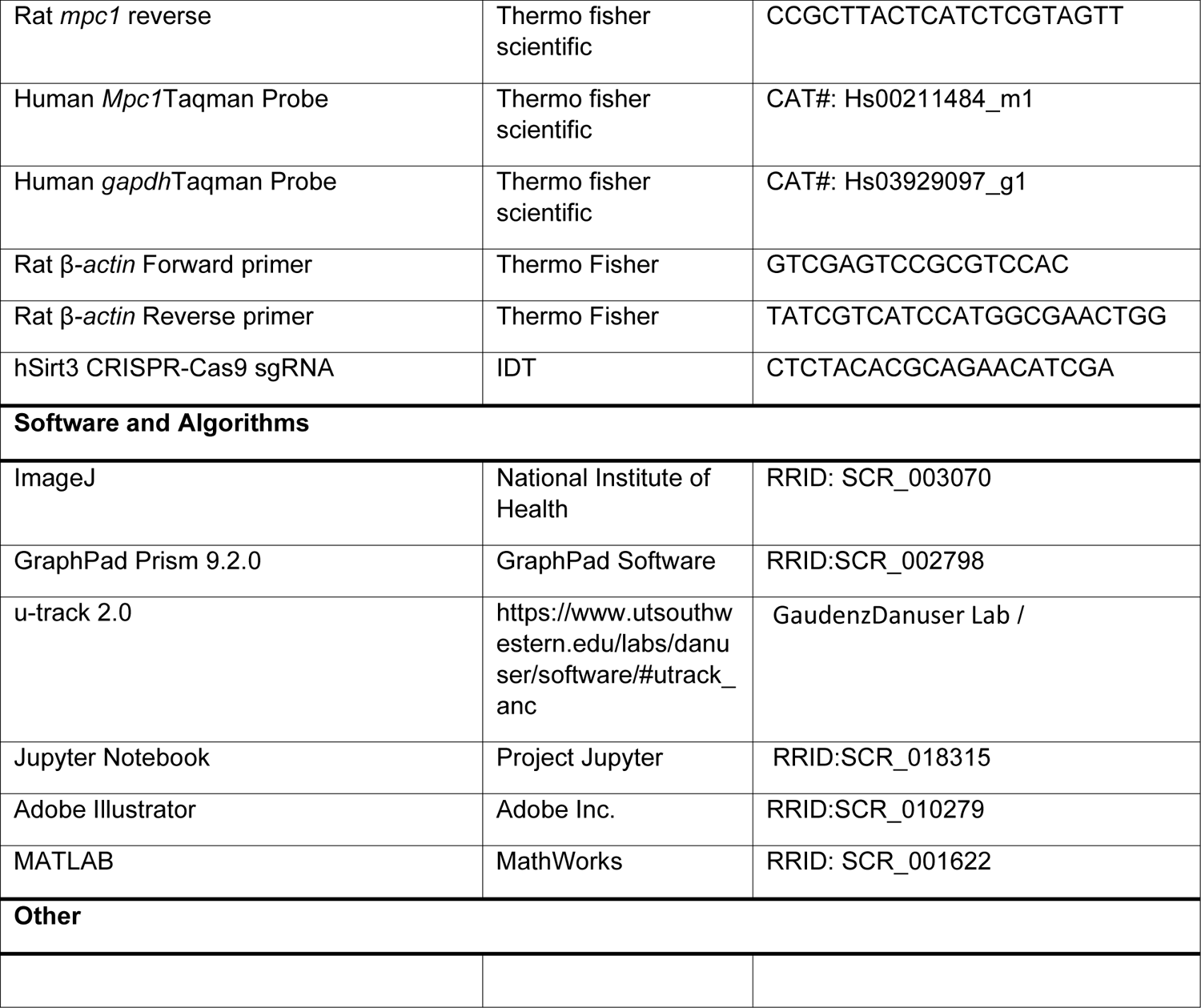

### STAR METHODS

#### Resource Availability

##### Lead Contact

Requests for further information and reagents should be directed to and will be fulfilled by the lead contact at.

##### Material Availability

Plasmids generated in this study will be available in the Addgene repository.

##### Data and Code Availability

Syn-ATP analysis code is available on GitHub under the directory AshrafiLab/Syn-ATP analysis.

### Experimental Model and Subject Details

#### Animals

Animal-related experiments were performed in accordance with protocols approved by the Washington university IACUC using wild-type rats of the Sprague-Dawley strain (RRID: RGD_734476), 129S6 wild-type and Sirt3^tm1.1Cxd^/ Sirt3^tm1.1Cxd^mice^33^, and C57BL/6for pyruvate tracing experiments.

#### Primary neuronal cultures

Hippocampi were dissected from 1–3-day old Sprague-Dawley rat pups(mixed-sex) and dissociated. Mixed cultures of neurons and glia were plated on coverslips coated with poly-ornithine and after 6–8 days of plating, were transfected with calcium phosphate, as previously described (Ryan, 1999, Tiwari et al., 2024). Hippocampal neurons were cultured in media containing 0.6% glucose, 0.1 g/L bovine transferrin (Millipore 616420), 0.25 g/L insulin, 0.3 g/L GlutaMAXTM supplement (Thermofisher 35050-061), 2% N-21 (R&D Systems AR008), 5% fetal bovine serum (R&D Systems S11510). 4 μm cytosine β-d-arabinofuranoside (Ara-C) was added at days in vitro (DIV) 2-3 to limit glial proliferation. Cortical neurons were maintained in culture media containing Neurobasal-A (NB-A) (Gibco # A24775-01) supplemented with 2% N-21, 5% fetal bovine serum, and other nutrients. Cultures were incubated at 37°C in a 95% air and 5% CO_2_ humidified incubator and experiments were performed on DIV 14 - 21 (hippocampal neurons) or DIV 10 - 14 (cortical neurons).

#### HEK293 and U2OS cell cultures

Cells were maintained in culture media composed of DMEM (Gibco11966-025), 10% FBS (Gibco 26140-079) and Pen/Strep (10,000 Units/mL) at 37°C in a 95% air and 5% CO_2_ humidified incubator. Cells were transfected at 30-40% confluency by calcium phosphate method or Lipofectamine 2000 (Thermofisher 11668019).

### Method Details

#### Plasmid Constructs

All the plasmid constructs and key reagents used in this paper are listed in the Key Resources Table. The following previously published DNA constructs were used: Syn-ATP^26^, vGLUT1-pHluorin^40^, and mito-PyronicSF (Addgene plasmid 124813)^23^. CaMKII-mito-pyronicSF construct was generated through insertion of mito-PyronicSF into the BamHI and HindIII sites of CaMKII-GCaMP6f.WPRE.SV40 (Addgene plasmid 100834). Wildtype shRNA-resistant (shR) MPC1 construct was generated by introducing 3 silent mutations in the regions complementary to shRNA in the mouse MPC1 sequence fused to a C-terminal FLAG tag. Wildtype shR MPC1 and MPC1-R_45_R_46_ were codon optimized for rat expression using the GeneOptimizer tool (ThermoFisher), synthesized (GeneArt Gene Synthesis, ThermoFisher) and cloned into the BamHI and EcoRI sites of pcDNA3.0. We used the pLKO.1 (Addgene plasmid 10878)^57^and pLKO.1-TRCmTagBFP2 (Addgene plasmid 191566) vectors for expression of shRNAsagainst *mpc1*(rat target sequence: CAAACGAAGTCGCTCAGCTCA, human target sequence: GCCTCGGAACTGGCTTCTGT) and*sirt3* (human target sequence: human target: GTGGGTGCTTCAAGTGTTGTT).

#### MTT survival assay

Cells were plated at a density of 1×10^5^ cells/well in pre-coated 96 well-plates in NB-A feeding medium. At DIV 8, medium was replaced for with media containing either glucose (5mM) or mixture of lactate and pyruvate (5mM each), or starvation media (only NB-A) and placed in a 5%CO2 incubator at 37°C for 16h. MTT solution was then added to each well for 4hours followed by incubation with a solubilizing solution overnight. The optical density was then measured at 450nm according to the manufacturer’s instructions (Cell proliferation kit I [MTT], Roche 11465007001). The background (medium only) was subtracted from every well and the survival rate was normalized to the mean survival of the control wells in 5mM Glucose.

#### Production and application of lentiviral particles

Lentiviral particles were produced by The Hope Center Viral Vectors Core at Washington University School of Medicine. BFP expression driven by CMV promoter on pLKO.1-TRC mTagBFP2 was used to establish the precise volume of lentivirus needed to induce maximal transduction of hippocampus or cortical cultures. Lentivirus was added on DIV 5 and media exchange was done after 2 days of infection. All experiments were carried out after 7 days of viral transduction as described previously^29^.

#### RNA isolation and quantitative PCR

RNeasy Mini Kits (QIAGEN 74104) were used for total RNA extraction from primary dissociated cortical neuron cultures. Reverse transcription was performed to generate cDNA using an All-In-One 5X RT Master Mix (ABM-G592) following the manufacturer’s instructions. Using cDNA as the template and either TaqMan® Fast Advanced Master Mix (Applied Biosytems, 4444557) or PowerUpTM SYBRTM Green Master Mix (Thermo Fisher, A25742), qPCR was set up. The ΔΔCt method was utilized to normalize relative mRNA expression to housekeeping genes, namely Gapdhor β-actin.

#### Western blotting

Neuronal extraction buffer (87792, Thermo Fisher Scientific) mixed with 1X protease inhibitor cocktail and 0.1 mM PMSF was used to solubilize cultured cortical cells and brain tissues. For protein quantification, BCA assay kit (BioVision), BioTek Synergy plate reader and Gen5 software (BioTek Instruments, Winooski, VT) were used. Immunoblotting was conducted using a 12 % SDS gradient polyacrylamide gel followed by transfer on PVDF membrane. The primary antibodies used in this paper are as follows: ATPase5β (1:1000, Sigma-Aldrich HPA001520), β-actin (1:5000, BioRad HCA147P), α-tubulin (1:2000, Sigma-Aldrich T6074), Sirt3 (1:1000, Cell Signaling Technology 5490S), and anti-acetyl lysine (1:1000, Cell Signaling Technology 9441S).

#### Immunoprecipitation of MPC1 from mouse brain

Cortical tissues from 5 *Sirt3^+/+^* and 5 *Sirt3^-/-^*mouse (6-10 weeks old, mixed sex) were homogenized in immunoprecipitation (IP) buffer (15 mM NaCl, 25 mM Tris Base, 1 mM EDTA (0.19 g), 0.2% NP-40 (1 mL), 10% Glycerol, 0.2 mM NaVO3, 0.5mM NaF, 0.1mM PMSF and 1X protease inhibitor cocktail), and sonicated. Homogenates were then centrifuged for 30 minutes at 4°C., and the supernatant was pre-cleared using Protein A agarose beads (30µl) for 2 hours at 4°C. The beads were spun down and the supernatant was incubated with normal rabbit IgG (Millipore 12-370; 3µg) and MPC1 antibody (gift of Dr. Brian Finck; 3µg) overnight at 4°C. Protein A agarose beads (30µl) were then added and incubated for 4 hours at 4°C. The supernatant was collected as unbound fraction after centrifugation and the beads were boiled at 100°C for 10 min in elution buffer (IP and 1x Lamelli buffer).

#### Generation of Sirt3 KD HEK293 cells using CRISPR-Cas9 gene editing

HEK293 cells were trypsinized and washed with PBS. The ribonucleoprotein (RNP) complex was prepared using sgRNA against Sirt3 (IDT), Cas 9 protein (Alt-R™S.p. Cas9 Nuclease V3, #1081058, IDT) and nucleofection kit SF (# V4XC-2012, Lonza) as per the manufacturer’s protocol. Program DG-130 was used on 4D-Nucleofactor system (Lonza) for transfection.

#### Immunoprecipitation of MPC1 from Sirt3 KD HEK293 cells

Sirt3 KD HEK293 cells generated with CRISPR/Cas9 (see above) or expressing shRNA against *sirt3* were transfected with either MPC1 WT-FLAG or MPC1 RR-FLAG construct using lipofectamine 2000 (#11668019,Invitrogen). After 24 hours of transfection, cells were harvested and lysed in IP buffer buffer (15 mM NaCl, 25 mM Tris Base, 1 mM EDTA (0.19 g), 0.2% NP-40 (1 mL), 10% Glycerol, 0.2 mM NaVO3, 0.5mM NaF, 0.1mM PMSF and 1X protease inhibitor cocktail). The supernatant was collected after centrifugation at 13000rpm for 20 mins and incubated with anti-FLAG conjugated magnetic beads. The beads were then pulled down by magnetic rack and washed thrice with the IP buffer. The protein was then eluted from the beads by boiling them in elution buffer (IP and 1x Lamelli buffer) at 100°C for 10 min.

#### Immunofluorescence and Confocal Microscopy

PFA (4%) was used for fixation of primary Hippocampal neurons. Neurons were then permeabilized with 0.5 % Triton and blocked with 5% bovine serum albumin (BSA) for 1 hour at room temperature (RT), and incubated at room temperature for 2 hours or overnight at 4°C with the following primary antibodies: MPC1 antibody (1:500, Sigma HPA045119) guinea pig vGLUT1 (1:500, Sigma AB5905); TOMM20 (1:500, Cell Signaling Technology 42406S). Coverslips were then incubated with the following secondary antibodies: anti-rabbit Alexa-fluor568 (1:500, Thermo FisherA21428) and anti-mouse Alexa-fluor488 (1:500, Thermo Fisher A11059) for 1 hour at RT and mounted with anti-fade mounting media (Thermo Fisher P36965), and kept at 4°C until imaged. Confocal images were collected on a Zeiss LSM 880 Confocal Microscope at the Washington University Center for Cellular Imaging which was purchased with support from the Office of Research Infrastructure Programs (ORIP), as part of the NIH Office of the Director under grant OD021629.

#### Live imaging of neurons and HEK293 cells

A custom-built laser-illuminated epifluorescence microscope with a U Plan Fluorite 40X Oil Objective (NA 1.30) and an Andor iXon Ultra 897camera cooled to −80°C to −95°C was used for live imaging experiments. Coverslips weremounted up in a laminar flow perfusion chamber and perfused with Tyrodes buffer consisting of (in mM) 119 NaCl, 2.5 KCl, 2 CaCl_2_, 2 MgCl_2_, 50 HEPES (pH 7.4), 5 glucose or 1.25 lactate and 1.25 pyruvate, supplemented with 10 μM 6-cyano-7nitroquinoxalibe-2, 3-dione (CNQX), and 50 μM D,L-2-amino-5phosphonovaleric acid (APV) (both from Sigma-Aldrich) to inhibit post-synaptic responses in neurons (these were not applied to HEK293 experiments). In experiments using neurons, action potentials were triggered using platinum-iridium electrodes with 1 ms pulses, producing field potentials of around 10 V/cm. All imaging was performed at 37°C using an Okolab stage top incubator for temperature control. As a standard, 30 frames were recorded before the stimulus train was triggered.

#### Near-TIRF microscopy of hippocampal synapses

All experiments were conducted at 37°C within a whole-microscope incubator chamber (TOKAI HIT). Fluorophores were excited with a 488 laser (Cell CMR-LAS-488, Olympus), and monitored using an inverted TIRF-equipped microscope (IX83, Olympus) under a 150x/1.45NA objective (UapoN). The Z-drift compensation system (IX3-ZDC) was used to ensure a constant position of the focal plane during imaging. Near-TIRF was achieved by adjusting the incident angle to 63.7°, which is near the critical angle of 63.63°. Images were acquired every 50 ms using a cooled EMCCD camera (iXon life 888, ANDOR). Field simulation was performed by using a pair of platinum electrodes and controlled by the software via Master-9 stimulus generator (A.M.P.I.).

#### Live imaging of mitochondrial pyruvate uptake

HEK293 cells were transfected with a fluorescence-based pyruvate sensor, mt Pyronic SF^23^ only (control) or together with shRNA against *sirt3* (Sirt3 KD) or *mpc1* (MPC1 KD). Cells were first imaged in the same Tyrodes buffer as described above but without <u>CNQX</u> and APV. After recording 50 frames, cells were perfused with Tyrodes supplemented with 10mM pyruvate. Once the fluorescence intensity reached a maximal plateau, cells were reperfused with 0 mM pyruvate. Areas of the coverslip typically containing 5-10 transfected cells were randomly selected for imaging.

#### TMRM Staining of HEK293 Cells

HEK cells were transfected either with PAAV-GFP construct only (control) or together with shRNA against *sirt3* (Sirt3KD) or *mpc1*(MPC1 KD) using Lipofectamine 2000 (ThermoFisher Scientific) and incubated for 45hr at 37°C in a 95% air/5% CO_2_ humidified incubator. Cells were then washed twice with MEM (Thermo Fisher Scientific) and stained with 25nM TMRM for 20mins at 37°C. Cells were washed thrice with MEM, incubated in 5mM TMRM, and imaged on the same custom-built laser-illuminated epifluorescence microscope as described above. GFP transfected cells were selected for analysis and the intensity of TMRM was measured using ImageJ, normalized to the mean of control. and plotted using GraphPad prism v9.0.

#### Analysis of live imaging data

Image analysis was primarily performed using the ImageJ plugin Time Series Analyzer where ∼20-50 regions of interest (ROIs) of ∼2 μm corresponding to responding nerve terminals were selected and fluorescence intensity was measured over time.HEK293 cells were analyzed by drawing ROIs corresponding to the entire cell volume.

#### Quantification of presynaptic ATP level

Dual luminescence and fluorescence imaging of the presynaptic ATP reporter, Syn-ATP, was performed using a semi-automatic platform as previously reported^29,58^. pH correction of luminescence/fluorescence (L/F) values was performed by collecting parallel measurements using the cytosolic pH sensor cyto-pHluorin expressed in neurons, as previously described^26^.

#### Quantification of SV retrieval block

Using, SV retrieval in hippocampus neurons expressing vGLUT1pHluorin was quantified as previously described^46^. Briefly, Images were captured at a frame rate of 2 Hz, and neurons were electrically stimulated with a train of 100 AP at 10 Hz. For experiments with MPC inhibitors, neurons were incubated with 100uM UK5099 or Zaprinast for 7min before imaging. Endocytic time constants (τ) were determined by fitting fluorescence change (ΔF) after the stimulation to a single exponential decay^59^. The proportion of ΔF left after twice the average endocytic time constant of the control (2τ), was divided by maximal ΔF at the end of stimulation (i.e. ΔF_2τ_/ΔF_max_), was used to compute the fractional retrieval block in the endocytosis of vGLUT1-pH.

#### Quantification of release probability

Release sites were defined using a hierarchical clustering algorithm using built-in functions in Matlab as described^41^. We have previously shown that the observed clusters do not arise from random distribution of release events, but rather represent a set of defined and repeatedly reused release sites within the AZs.

#### Event detection and localization using mixture-model fitting

The release event detection and localization at subpixel resolution were performed as previously described^41^ using MATLAB and the uTrack software package, which was kindly made available by Dr Gaudenz Danuser’s lab ^60,61^. Localization precision was determined directly from least-squares Gaussian fits of individual events as previously described^43^.

#### Definition of AZ dimensions and center

The AZ size was approximated based on the convex hull encompassing all vesicle fusion events in a given bouton. This measurement is in close agreement with the ultrastructural measurements of AZ dimensions^41^. AZ center was defined as the mean position of all fusion events in a given bouton.

#### Metabolic profiling of cultured cortical neurons

Cortical neurons were plated at a density of 1×10^6^ cells/well in pre-coated 6-well plates. At DIV 7, medium was replaced with medium containing either glucose (5mM) or a mixture of lactate and pyruvate (5mM each). Nutrient uptake was determined by the method described before^62, 63^ with minor modifications. Media were collected after cells were incubated for 24 h. U-^13^C labeled nutrients (glucose, lactate, glutamine, and glutamate; Cambridge Isotope Laboratories) were added into the media as the internal standards. Media extraction and metabolite detection were conducted accordingly. The quantification was performed by calculating the ratio between the labeled peak and the unlabeled peak of the same metabolite in each media sample. The nutrient consumption rate was finally normalized by the cell numbers and proliferation rate.

#### Jugular vein catheterization

To perform infusion studies, a catheter (Instech, C20PU-MJV1301) was placed in the right jugular vein and connected to a vascular access button (Instech, VABM1B/25) implanted subcutaneously in the back of the mice. All catheter implantation surgeries were performed at the Hope Center for Neurological Diseases, Washington University. Mice were allowed to recover from surgery for at least one week before tracer infusion.

#### Intravenous infusion of mice with U^13^C-Pyruvatefor metabolomics analysis

On the day of the procedure, U^13^C-Pyruvate (CIL, CLM-2440-PK) was freshly prepared in saline at a concentration of 400mM. The mice were weighed to calculate the tracer infusion rate and fasted at 9 am (ZT2). At approximately 2:00 pm, the vascular access button of individual mice was connected to the infusion line with a swivel (Instech, SMCLA), tether (Instech, KVABM1T/25), and infusion pump (CHEMYX, Fusion 100T). The infusion line was prefilled with 400mM U^13^C-Pyruvate. Prime infusion was initiated at 1ul/min/g for 2 minutes, followed by continuous infusion at 0.1ul/min/g for 2 hours. Following completion of the pyruvate infusion, mice were anesthetized, and blood was collected by cardiac puncture. Tissues were subsequently collected as quickly as possible (in 10 min or less) following euthanasia and snap-frozen in liquid nitrogen. Tissues were stored at −80 °C until processing for LC/MS analysis.

#### Metabolite extraction from serum and tissues

To extract metabolites from serum, 5ul serum was mixed with 295ul ice-cold methanol:acetonitrile:water (2:2:1), vortexed, and incubated at −20 °C for 1 h. The liver and brain tissues were ground by pestle and mortar with liquid nitrogen. Ground tissue mixed with ice-cold methanol:acetonitrile:water (2:2:1), and subjected to two cycles of 7m/s (30 s/cycle) using an Omni Bead Ruptor Elute Homogenizer. For every 1 mg of tissue wet weight, 30 μL of extraction solvent was added. Samples were then incubated at −20 °C for 1 h to precipitate protein. Serum and Tissue extracts were centrifuged at 20,000 g and 4 °C for 10 min, and the supernatant was transferred into LC/MS vials.

#### Metabolite measurement with LC-MS

Ultra-high-performance LC (UHPLC)/MS was performed with a Thermo Scientific Vanquish Horizon UHPLC system interfaced with a Thermo Scientific Q Exactive Plus Orbitrap mass spectrometer. Metabolites were separated on a HILICON iHILIC-(P)-Classic column (100 x 2.1 mm, 5 μm). The injection volume was 5 μL. The column compartment was maintained at 40 °C. The mobile-phase solvents were composed of: A = 20 mM ammonium bicarbonate, 2.5 μM medronic acid, 0.1% ammonium hydroxide in water:acetonitrile 95:5; and B = water:acetonitrile 5:95. The following linear gradient was applied at a flow rate of 250uL min^-1^: 0 – 1min, 90% B; 12min, 35% B; 12.5 – 14.5min, 25% B; 15min, 90% B followed by a re-equilibration phase of 10 column volumes. Data was acquired in negative ion mode with the following settings: spray voltage, 2.8 kV; sheath gas, 45 Arb; auxiliary gas, 10 Arb; sweep gas, 2 Arb; capillary temperature, 250 °C; aux gas temperature, 350 °C; mass range, 65 – 975 Da; resolution, 140,000, automatic gain control (AGC) target 1e6, maximum injection time 200 milliseconds (ms). LC/MS data were processed and analyzed with the open-source Skyline software^64^. Naturally occurring isotope distributions were corrected as previously described^65^. Metabolite levels were plotted with GraphPad Prism. To correct for multiple comparison, raw P values were adjusted using the FDR approach using the two-stage step-up method of Benjamini, Krieger, and Yekutielii.An FDR adjusted p value <0.01 was considered statistically significant.

#### Statistical Analysis

GraphPad Prism v10.0 or MATLAB were used for statistical analysis. When comparing two sets of data, P values were determined using the two-tailed, unpaired t test, or the nonparametric Mann-Whitney U test. For more than two datasets, two-way ANOVA test was employed. A P value of less than 0.05 was deemed statistically significant. Based on power analysis and estimations of data variability derived from earlier findings and pilot research, the appropriate sample size was chosen. Grubb’s test in GraphPad Prism was used to identify and eliminate outliers from further analysis.

## SUPPLEMENTARY FIGURE LEGENDS

**Figure S1 (Related to** Figure 2**). Quantification of neuronal MPC1 knockdown with qPCR.** Relative expression of *mpc1* mRNA in control and MPC1 KD cortical neurons. Values are normalized to *β-actin* mRNA and expressed relative to the control. n= 3 cortical cultures. Mann-Whitney U test.

**Figure S2** (Related to Figure 3). Quantification of Sirt3 and MPC1 knockdown efficiency in HEK293 cells and characterization of mitochondrial membrane potential. **(A)** Relative *mpc1*mRNA expression in control and MPC1 KD HEK293 cells. Values are normalized to *gapdh* mRNA and expressed relative to control. n =9 lysates. Average normalized mRNA level ± SEM: Control, 1.0 ± 0.0; *mpc1* KD, 0.4 ± 0.06. **(B)** Total protein lysate from control HEK293 cells and cells expressing shRNA against Sirt3 were immunoblotted for Sirt3 and mitochondrial ATPase5β. **(C)** Relative expression of Sirt3 quantified from blots shown in panel E. Values are normalized to ATPase5β and expressed relative to the control. n = 6 (lysates/blots). **(D)** Fluorescence intensity of TMRM staining in MPC1 and Sirt3 KD cells normalized to control (untransfected) cells. n = 318-388 (cells). **(E)** Relative expression of Sirt3 protein in control HEK293 cells or Sirt3 KD cells created with CRISPR-Cas9 editing of *sirt3* gene, probed across successive culture passages (denoted as P). Immunoblotting against mitochondrial ATPase5β was used as loading control. n=1 western blot, 6 total lysates. Mann-Whitney U test (A and C), Kruskal-Wallis test (D). *p < 0.05; ** p < 0.01, ***p < 0.001.

**Figure S3** (Related to Figure 4). Pharmacological and genetic MPC impair SV retrieval in hippocampal nerve terminals. **(A, C)** Sample normalized vGLUT1-pH traces in hippocampal terminals electrically stimulated with 100 AP at 10 Hz before (control) and after treatment with MPC inhibitors UK5099 (A), or Zaprinast (C). **(B, D)** SV retrieval quantified as fractional retrieval block from traces in A and C. **(B)** n = 9 (neurons). **(D)** n = 10 (neurons). **(E)** Sample normalized vGLUT1-pH traces in hippocampal terminals during stimulation with 100 AP at 10 Hz in control, MPC1 KD neurons or KD neurons expressing shRNA-resistant MPC1 (MPC1-wt). **(F)** SV retrieval quantified as fractional retrieval block from traces in E. n = 18-81(neurons). Black bar denotes electrical stimulation. Kruskal-Wallis test (B and D), Mann-Whitney U test (F). *p < 0.05.

**Tables S1 (Related to** Figure 1**). Dataset of Isotopologues derived from ^13^C_3_ pyruvate tracing in mouse serum and tissues.**

**Tables S2 (Related to** Figure 1**). Metabolite pool sizes in primary cortical neurons supplied with glucose or lactate/pyruvate.**

**Table S3 (Related to** Figures 1-4**, and S1-3). Statistical data for figures.**

## Notes

### Competing Interest Statement

The authors have declared no competing interest.

