## Supplementary Figures for "Mitochondrial pyruvate transport regulates presynaptic metabolism and neurotransmission"

A

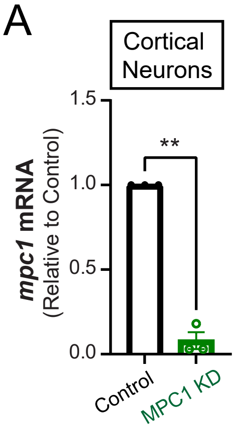

Cortical Neurons
1.5
1.0
0.5
0.0
**
mpc1 mRNA
(Relative to Control)
Control
MPC1 KD


Figure S1

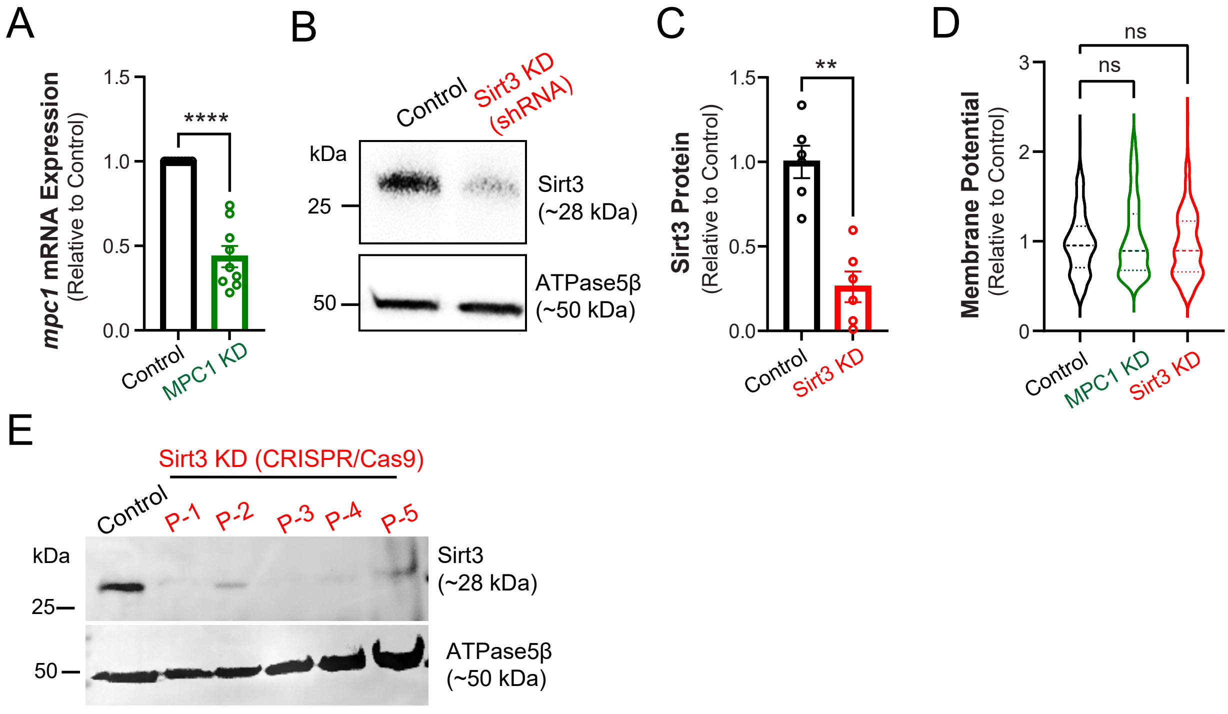
A
mpc1 mRNA Expression
(Relative to Control)
1.5
1.0
0.5
0.0
****
Control
MPC1 KD
B
Control
Sirt3 KD (shRNA)
kDa
25
50
Sirt3 (~28 kDa)
ATPase5β (~50 kDa)
C
Sirt3 Protein
(Relative to Control)
1.5
1.0
0.5
0.0
**
Control
Sirt3 KD
D
Membrane Potential
(Relative to Control)
3
2
1
0
ns
ns
Control
MPC1 KD
Sirt3 KD
E
Sirt3 KD (CRISPR/Cas9)
Control
P-1
P-2
P-3
P-4
P-5
kDa
25
50
Sirt3 (~28 kDa)
ATPase5β (~50 kDa)

Figure S2

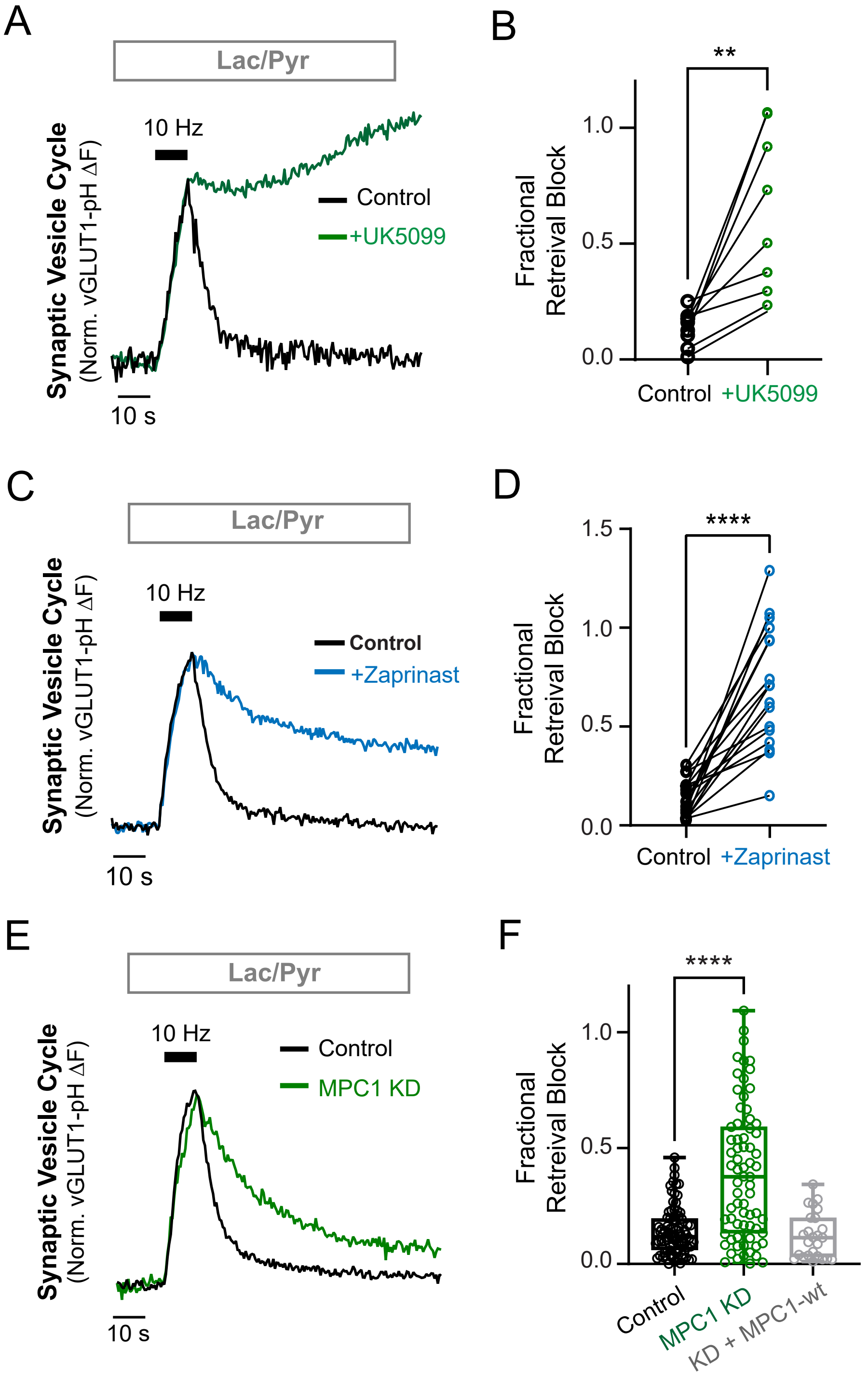
A
Lac/Pyr
10 Hz
Control
+UK5099
Synaptic Vesicle Cycle
(Norm. vGLUT1-pH ΔF)
10 s
B
**
1.0
0.5
0.0
Fractional
Retreival Block
Control
+UK5099
C
Lac/Pyr
10 Hz
Control
+Zaprinast
Synaptic Vesicle Cycle
(Norm. vGLUT1-pH ΔF)
10 s
D
****
1.5
1.0
0.5
0.0
Fractional
Retreival Block
Control
+Zaprinast
E
Lac/Pyr
10 Hz
Control
MPC1 KD
Synaptic Vesicle Cycle
(Norm. vGLUT1-pH ΔF)
10 s
F
****
1.0
0.5
0.0
Fractional
Retreival Block
Control
MPC1 KD
KD + MPC1-wt

Figure S3

| Isotopologues | Pyruvate | | | | | Lactate | | | | |
|---|---|---|---|---|---|---|---|---|---|---|
| | [m+0] | [m+1] | [m+2] | [m+3] | Total label | [m+0] | [m+1] | [m+2] | [m+3] | Total label |
| Serum_A | 0.93 | 0.02 | 0.02 | 0.03 | 0.07 | 0.93 | 0.02 | 0.02 | 0.03 | 0.07 |
| Serum_B | 0.91 | 0.02 | 0.02 | 0.06 | 0.09 | 0.91 | 0.02 | 0.02 | 0.06 | 0.09 |
| Serum_C | 0.92 | 0.02 | 0.02 | 0.05 | 0.08 | 0.92 | 0.02 | 0.02 | 0.05 | 0.08 |
| Serum_D | 0.93 | 0.02 | 0.02 | 0.03 | 0.07 | 0.93 | 0.02 | 0.02 | 0.03 | 0.07 |
| Brain_A | 0.95 | 0.02 | 0.02 | 0.01 | 0.05 | 0.95 | 0.02 | 0.02 | 0.01 | 0.05 |
| Brain_B | 0.97 | 0.01 | 0.01 | 0.01 | 0.03 | 0.95 | 0.02 | 0.02 | 0.01 | 0.05 |
| Brain_C | 0.95 | 0.02 | 0.01 | 0.02 | 0.05 | 0.94 | 0.02 | 0.02 | 0.02 | 0.06 |
| Brain_D | 0.95 | 0.02 | 0.02 | 0.02 | 0.05 | 0.94 | 0.02 | 0.02 | 0.02 | 0.06 |
| Liver_A | 0.98 | 0.01 | 0.01 | 0.00 | 0.02 | 0.97 | 0.01 | 0.01 | 0.01 | 0.03 |
| Liver_B | 0.97 | 0.01 | 0.01 | 0.01 | 0.03 | 0.97 | 0.01 | 0.01 | 0.01 | 0.03 |
| Liver_C | 0.96 | 0.01 | 0.01 | 0.01 | 0.04 | 0.95 | 0.02 | 0.01 | 0.01 | 0.05 |
| Liver_D | 0.97 | 0.01 | 0.01 | 0.01 | 0.03 | 0.98 | 0.01 | 0.01 | 0.01 | 0.02 |

| Isotopologues | Glucose | | | | | | | | Hexose phosphate | | | | | | | |
|---|---|---|---|---|---|---|---|---|---|---|---|---|---|---|---|---|
| | [m+0] | [m+1] | [m+2] | [m+3] | [m+4] | [m+5] | [m+6] | Total label | [m+0] | [m+1] | [m+2] | [m+3] | [m+4] | [m+5] | [m+6] | Total label |
| Serum_A | 0.89 | 0.05 | 0.04 | 0.03 | 0.00 | 0.00 | 0.00 | 0.11 | 0.85 | 0.00 | 0.02 | 0.00 | 0.14 | 0.00 | 0.00 | 0.15 |
| Serum_B | 0.90 | 0.04 | 0.03 | 0.03 | 0.00 | 0.00 | 0.00 | 0.10 | 0.78 | 0.00 | 0.00 | 0.00 | 0.21 | 0.00 | 0.01 | 0.22 |
| Serum_C | 0.87 | 0.06 | 0.04 | 0.03 | 0.00 | 0.00 | 0.00 | 0.13 | 0.58 | 0.00 | 0.00 | 0.05 | 0.37 | 0.00 | 0.00 | 0.42 |
| Serum_D | 0.89 | 0.04 | 0.04 | 0.03 | 0.00 | 0.00 | 0.00 | 0.11 | 0.78 | 0.00 | 0.00 | 0.00 | 0.21 | 0.00 | 0.00 | 0.22 |
| Brain_A | 0.95 | 0.00 | 0.05 | 0.00 | 0.00 | 0.00 | 0.00 | 0.05 | 1.00 | 0.00 | 0.00 | 0.00 | 0.00 | 0.00 | 0.00 | 0.00 |
| Brain_B | 0.96 | 0.00 | 0.01 | 0.00 | 0.00 | 0.00 | 0.02 | 0.04 | 0.99 | 0.00 | 0.00 | 0.00 | 0.00 | 0.00 | 0.00 | 0.01 |
| Brain_C | 0.99 | 0.00 | 0.01 | 0.00 | 0.00 | 0.00 | 0.00 | 0.01 | 0.99 | 0.00 | 0.01 | 0.00 | 0.00 | 0.00 | 0.00 | 0.01 |
| Brain_D | 0.99 | 0.00 | 0.01 | 0.00 | 0.00 | 0.00 | 0.00 | 0.01 | 0.99 | 0.00 | 0.00 | 0.00 | 0.00 | 0.00 | 0.00 | 0.01 |
| Liver_A | 0.97 | 0.02 | 0.01 | 0.01 | 0.00 | 0.00 | 0.00 | 0.03 | 0.99 | 0.00 | 0.01 | 0.00 | 0.00 | 0.00 | 0.00 | 0.01 |
| Liver_B | 0.97 | 0.02 | 0.01 | 0.01 | 0.00 | 0.00 | 0.00 | 0.03 | 0.99 | 0.00 | 0.01 | 0.00 | 0.00 | 0.00 | 0.00 | 0.01 |
| Liver_C | 0.89 | 0.05 | 0.03 | 0.02 | 0.00 | 0.00 | 0.00 | 0.11 | 0.99 | 0.00 | 0.01 | 0.00 | 0.00 | 0.00 | 0.00 | 0.01 |
| Liver_D | 0.97 | 0.01 | 0.01 | 0.01 | 0.00 | 0.00 | 0.00 | 0.03 | 0.99 | 0.00 | 0.01 | 0.00 | 0.00 | 0.00 | 0.00 | 0.01 |

| Isotopologues | Fructose Bis-phosphate | | | | | | | | Citrate | | | | | | | |
|---|---|---|---|---|---|---|---|---|---|---|---|---|---|---|---|---|
| | [m+0] | [m+1] | [m+2] | [m+3] | [m+4] | [m+5] | [m+6] | Total label | [m+0] | [m+1] | [m+2] | [m+3] | [m+4] | [m+5] | [m+6] | Total label |
| Serum_A | 0.72 | 0.00 | 0.00 | 0.00 | 0.08 | 0.12 | 0.08 | 0.28 | 0.92 | 0.05 | 0.02 | 0.01 | 0.00 | 0.00 | 0.00 | 0.08 |
| Serum_B | 0.48 | 0.05 | 0.07 | 0.00 | 0.21 | 0.13 | 0.06 | 0.52 | 0.94 | 0.03 | 0.01 | 0.01 | 0.00 | 0.00 | 0.00 | 0.06 |
| Serum_C | 0.06 | 0.00 | 0.14 | 0.00 | 0.04 | 0.00 | 0.77 | 0.94 | 0.94 | 0.04 | 0.02 | 0.01 | 0.00 | 0.00 | 0.00 | 0.06 |
| Serum_D | 0.37 | 0.00 | 0.13 | 0.00 | 0.22 | 0.12 | 0.16 | 0.63 | 0.93 | 0.04 | 0.02 | 0.01 | 0.00 | 0.00 | 0.00 | 0.07 |
| Brain_A | 0.98 | 0.00 | 0.02 | 0.00 | 0.00 | 0.00 | 0.00 | 0.02 | 0.83 | 0.11 | 0.06 | 0.01 | 0.00 | 0.00 | 0.00 | 0.17 |
| Brain_B | 0.98 | 0.00 | 0.02 | 0.00 | 0.00 | 0.00 | 0.00 | 0.02 | 0.85 | 0.10 | 0.04 | 0.00 | 0.00 | 0.00 | 0.00 | 0.15 |
| Brain_C | 0.98 | 0.00 | 0.02 | 0.00 | 0.00 | 0.00 | 0.00 | 0.02 | 0.84 | 0.10 | 0.05 | 0.01 | 0.00 | 0.00 | 0.00 | 0.16 |
| Brain_D | 0.98 | 0.00 | 0.02 | 0.00 | 0.00 | 0.00 | 0.00 | 0.02 | 0.84 | 0.10 | 0.05 | 0.01 | 0.00 | 0.00 | 0.00 | 0.16 |
| Liver_A | 0.94 | 0.00 | 0.02 | 0.01 | 0.02 | 0.00 | 0.00 | 0.06 | 0.93 | 0.05 | 0.01 | 0.01 | 0.00 | 0.00 | 0.00 | 0.07 |
| Liver_B | 0.96 | 0.00 | 0.02 | 0.01 | 0.01 | 0.00 | 0.00 | 0.04 | 0.93 | 0.04 | 0.02 | 0.01 | 0.00 | 0.00 | 0.00 | 0.07 |
| Liver_C | 0.93 | 0.01 | 0.03 | 0.02 | 0.01 | 0.00 | 0.00 | 0.07 | 0.94 | 0.04 | 0.02 | 0.01 | 0.00 | 0.00 | 0.00 | 0.06 |
| Liver_D | 0.93 | 0.00 | 0.02 | 0.02 | 0.03 | 0.00 | 0.00 | 0.07 | 0.94 | 0.04 | 0.01 | 0.00 | 0.00 | 0.00 | 0.00 | 0.06 |

| Isotopologues | Succinate | | | | | | Malate | | | | | |
|---|---|---|---|---|---|---|---|---|---|---|---|---|
| | [m+0] | [m+1] | [m+2] | [m+3] | [m+4] | Total label | [m+0] | [m+1] | [m+2] | [m+3] | [m+4] | Total label |
| Serum_A | 0.88 | 0.04 | 0.08 | 0.00 | 0.00 | 0.12 | 0.93 | 0.04 | 0.02 | 0.01 | 0.00 | 0.07 |
| Serum_B | 0.60 | 0.12 | 0.19 | 0.00 | 0.09 | 0.40 | 0.93 | 0.04 | 0.02 | 0.01 | 0.00 | 0.07 |
| Serum_C | 0.84 | 0.02 | 0.07 | 0.00 | 0.06 | 0.16 | 0.93 | 0.04 | 0.02 | 0.01 | 0.00 | 0.07 |
| Serum_D | 0.89 | 0.00 | 0.09 | 0.00 | 0.01 | 0.11 | 0.94 | 0.04 | 0.02 | 0.01 | 0.00 | 0.06 |
| Brain_A | 0.84 | 0.12 | 0.03 | 0.00 | 0.00 | 0.16 | 0.86 | 0.10 | 0.04 | 0.00 | 0.00 | 0.14 |
| Brain_B | 0.86 | 0.11 | 0.03 | 0.00 | 0.00 | 0.14 | 0.88 | 0.08 | 0.03 | 0.00 | 0.00 | 0.12 |
| Brain_C | 0.85 | 0.11 | 0.04 | 0.00 | 0.00 | 0.15 | 0.87 | 0.09 | 0.03 | 0.00 | 0.00 | 0.13 |
| Brain_D | 0.85 | 0.11 | 0.04 | 0.00 | 0.00 | 0.15 | 0.87 | 0.09 | 0.04 | 0.00 | 0.00 | 0.13 |
| Liver_A | 0.93 | 0.05 | 0.02 | 0.00 | 0.00 | 0.07 | 0.93 | 0.05 | 0.02 | 0.00 | 0.00 | 0.07 |
| Liver_B | 0.92 | 0.05 | 0.02 | 0.00 | 0.00 | 0.08 | 0.93 | 0.04 | 0.02 | 0.01 | 0.00 | 0.07 |
| Liver_C | 0.94 | 0.04 | 0.02 | 0.00 | 0.00 | 0.06 | 0.94 | 0.04 | 0.02 | 0.01 | 0.00 | 0.06 |
| Liver_D | 0.93 | 0.05 | 0.01 | 0.00 | 0.00 | 0.07 | 0.94 | 0.04 | 0.02 | 0.00 | 0.00 | 0.06 |

| Isotopologues | Glutamate | | | | | | | Aspartate | | | | | |
|---|---|---|---|---|---|---|---|---|---|---|---|---|---|
| | [m+0] | [m+1] | [m+2] | [m+3] | [m+4] | [m+5] | Total label | [m+0] | [m+1] | [m+2] | [m+3] | [m+4] | Total label |
| Serum_A | 0.93 | 0.04 | 0.03 | 0.00 | 0.00 | 0.00 | 0.07 | 0.96 | 0.03 | 0.01 | 0.00 | 0.00 | 0.04 |
| Serum_B | 0.94 | 0.03 | 0.02 | 0.00 | 0.00 | 0.00 | 0.06 | 0.95 | 0.03 | 0.01 | 0.01 | 0.00 | 0.05 |
| Serum_C | 0.95 | 0.03 | 0.02 | 0.00 | 0.00 | 0.00 | 0.05 | 0.97 | 0.02 | 0.01 | 0.00 | 0.00 | 0.03 |
| Serum_D | 0.94 | 0.04 | 0.02 | 0.00 | 0.00 | 0.00 | 0.06 | 0.95 | 0.03 | 0.01 | 0.00 | 0.00 | 0.05 |
| Brain_A | 0.84 | 0.10 | 0.06 | 0.00 | 0.00 | 0.00 | 0.16 | 0.88 | 0.08 | 0.04 | 0.00 | 0.00 | 0.12 |
| Brain_B | 0.86 | 0.09 | 0.05 | 0.00 | 0.00 | 0.00 | 0.14 | 0.89 | 0.08 | 0.03 | 0.00 | 0.00 | 0.11 |
| Brain_C | 0.85 | 0.10 | 0.05 | 0.00 | 0.00 | 0.00 | 0.15 | 0.88 | 0.08 | 0.03 | 0.00 | 0.00 | 0.12 |
| Brain_D | 0.84 | 0.10 | 0.05 | 0.00 | 0.00 | 0.00 | 0.16 | 0.88 | 0.08 | 0.03 | 0.00 | 0.00 | 0.12 |
| Liver_A | 0.95 | 0.03 | 0.02 | 0.00 | 0.00 | 0.00 | 0.05 | 0.95 | 0.04 | 0.01 | 0.00 | 0.00 | 0.05 |
| Liver_B | 0.95 | 0.02 | 0.02 | 0.00 | 0.00 | 0.00 | 0.05 | 0.95 | 0.03 | 0.02 | 0.00 | 0.00 | 0.05 |
| Liver_C | 0.96 | 0.02 | 0.02 | 0.00 | 0.00 | 0.00 | 0.04 | 0.95 | 0.03 | 0.01 | 0.00 | 0.00 | 0.05 |
| Liver_D | 0.96 | 0.02 | 0.02 | 0.00 | 0.00 | 0.00 | 0.04 | 0.96 | 0.03 | 0.01 | 0.00 | 0.00 | 0.04 |

Table S1

**Pyruvate Consumption Rate (Fig 1, F)**

| | | Ctrl media | | Neuronal culture | |
|---|---|---|---|---|---|
| Pyruvate Isotopologue | | 87.0088 [m+0] | 90.018865 [m+3] | 87.0088 [m+0] | 90.018865 [m+3] |
| +Glucose | sample 1 | 21488957 | 16786251105 | 239244145 | 14888723009 |
| | sample 2 | 25007884 | 15988839384 | 245954945 | 16859657350 |
| | sample 3 | 19099959 | 15173338453 | 226370605 | 15378550766 |
| +Lac/Pyr | sample 1 | 6878834864 | 15665260149 | 2321945972 | 15446645198 |
| | sample 2 | 6169316360 | 14445753724 | 2213495973 | 15171144997 |
| | sample 3 | 6226607930 | 14961289632 | 2444021211 | 16207574533 |

**Metabolite Levels (Fig 1, G)**

| | | Glucose | Pyruvate | Lactate | Alanine | Citrate | a-ketoglutarate | Succinate | Fumarate | Malate | Aspartate | Glutamate | Glutamine | Proline |
|---|---|---|---|---|---|---|---|---|---|---|---|---|---|---|
| +Glucose | sample 1 | 2.80E+07 | 2.81E+08 | 1.74E+11 | 1.89E+09 | 1.01E+10 | 1.36E+08 | 1.57E+09 | 6.99E+08 | 1.12E+10 | 4.39E+09 | 1.51E+10 | 8.54E+09 | 2.83E+08 |
| | sample 2 | 3.01E+07 | 2.80E+08 | 1.42E+11 | 1.91E+09 | 1.00E+10 | 1.23E+08 | 1.44E+09 | 4.26E+08 | 5.99E+09 | 3.93E+09 | 1.42E+10 | 7.74E+09 | 2.69E+08 |
| | sample 3 | 1.79E+07 | 2.01E+08 | 1.04E+11 | 1.78E+09 | 8.87E+09 | 1.24E+08 | 1.22E+09 | 3.67E+08 | 5.19E+09 | 4.00E+09 | 1.39E+10 | 6.88E+09 | 2.67E+08 |
| +Lac/Pyr | sample 1 | 3.62E+08 | 1.60E+09 | 1.02E+11 | 3.04E+09 | 1.30E+10 | 1.93E+09 | 1.28E+10 | 6.40E+08 | 9.52E+09 | 3.06E+09 | 1.99E+10 | 6.14E+09 | 3.80E+08 |
| | sample 2 | 2.65E+07 | 1.84E+09 | 1.64E+11 | 2.54E+09 | 1.39E+10 | 6.82E+09 | 8.66E+09 | 7.60E+08 | 1.07E+10 | 3.25E+09 | 2.19E+10 | 6.84E+09 | 3.90E+08 |
| | sample 3 | 9.02E+07 | 1.59E+09 | 1.08E+11 | 2.54E+09 | 1.25E+10 | 5.14E+08 | 9.69E+09 | 5.69E+08 | 8.39E+09 | 4.39E+09 | 1.72E+10 | 5.60E+09 | 3.19E+08 |

### Table S2

**Table S3: Statistical data for Figures**

| **Figure 1** | **Sample number** | **Mean ± SEM** | **Statistical test** |
|---|---|---|---|
| Figure 1C | 4 mice | see Supplementary Table 1 | Unpaired t-test |
| Figure 1E | 3 (culture); Glucose: 105 (wells), no Glucose: 108 (wells), Lac/Pyr: 107 (wells) | Glucose: 100 ± 1.5 ; no Glucose: 25.1 ± 1.4 ; Lac/Pyr: 57.6 ± 1.3 | One-Way ANOVA |
| Figure 1F | 1 (culture); Glucose: 3 (wells); Lac/Pyr: 3 (wells) | Glucose: -0.00007746 ± 9.943e-006; Lac/Py: 0.001423 ± 0.0001220 | Unpaired t-test |
| Figure 1G | 1 (culture); Glucose: 3 (wells); Lac/Pyr: 3 (wells) | Glucose: Pyruvate (1.00 ± 0.15), Citrate (1.00 ± 0.04), Malate (1.00 ± 0.25), Succincate (1.00 ± 0.07); Lac/Pyr: Pyruvate (6.60 ± 0.31), Citrate (1.36 ± 0.04), Malate (1.28 ± 0.09), Succincate (7.34 ± 0.87) | Multiple unpaired t-tests |
| **Figure 2** | **Sample number** | **Mean ± SEM** | **Statistical test** |
| Figure 2F,G | 3 (culture), Ctrl: 18 (neurons/coverslips), MPC1 KD: 12 (neurons/coverslips) | Ctrl: 0.72 ± 0.11; MPC1 KD: -0.01 ± 0.06 | Mann-Whitney U Test |
| Figure 2H,I | 3 (culture), 9 (neurons/coverslips) | 6-8 min with UK5099: 0.6 ± 0.11 | One sample t-test |
| **Figure 3** | **Sample number** | **Mean ± SEM** | **Statistical test** |
| Figure 3A, B | 5-27 (culture), Ctrl: 50 (coverslips), 57 (FOV); MPC1 KD: 24 (coverslips)/ 33 (FOV); Sirt3 KD: 10 (coverslips)/ 18 (FOV), #5-6 cells/ FOV | Ctrl: 0.4 ± 0.02; MPC1 KD: 0.03 ± 0.02; Sirt3 KD: 0.21 ± 0.3 | Kruskal-Wallis Test |
| Figure 3C, D | 4 (IP/blots), Sirt3$^{+/+}$: 20 (mouse); Sirt3$^{-/-}$: 20 (mouse) | Ac-K intensity norm. to MPC1 intenstiy: Sirt3$^{+/+,}$ 1.35 ± 0.44; Sirt3 $^{-/-,}$ 2.10 ± 0.64. Ac-K intensity norm. to control: Sirt3$^{-/-,}$ 1.6 ± 0.1 | Paired t-test of Ac-K intensity norm. to MPC1 intenstiy |
| Figure 3F, G | 6 (IP/blots) | Ac-K intensity norm. to MPC1-WT: MPC1 WT, 1.0 ± 0.0; MPC1 RR, 0.36 ± 0.08 | One-Sample t-test |
| **Figure 3** | **Sample number** | **Mean ± SEM** | **Statistical test** |
| Figure 4B | 3-9 (culture), Ctrl: 9 (Cultures) / 36 (Coverslips) / 1,163 (Syn); +UK5099: 3 (Cultures) / 9 (Coverslips) / 230 (Syn) | Ctrl: 0.0746 ± 0.001; +UK5099: 0.0649 ± 0.001 | Two-Sample t-test |
| Figure 4C | 3-9 (culture), Ctrl: 9 (Cultures) / 36 (Coverslips) / 1,254 (Syn); +UK5099: 3 (Cultures) / 9 (Coverslips) / 220 (Syn) | Ctrl: 9.25 ± 0.06; +UK5099: 6.30 ± 0.96 | Two-Sample t-test |
| Figure 4D | 3-9 (culture), Ctrl: 9 (Cultures) / 36 (Coverslips) / 11,925 (Release site); +UK5099: 3 (Cultures) / 9 (Coverslips) / 1,705(Release Site) | Ctrl: 1.462 ± 0.009; +UK5099: 1.634 ± 0.0239 | Two-Sample t-test |
| Figure 4E | 3-9 (culture), Ctrl: 9 (Cultures) / 36 (Coverslips) / 15,084 (events); +UK5099: 3 (Cultures) / 9 (Coverslips) / 2,786 events) | Ctrl: 177.281 ± 0.338 ;+UK5099: 146.608 ± 0.873 | Two-Sample t-test |
| Figure 4F | 3-9 (culture), Ctrl: 9 (Cultures) / 36 (Coverslips) / 15,084 (events); +UK5099: 3 (Cultures) / 9 (Coverslips) / 2,786 events) | Ctrl: 121.218 ± 0.623; +UK5099: 97.967 ± 1.415 | Two-Sample t-test |
| Figure 4H | 3-9 (culture), Ctrl: 9 (Cultures) / 36 (Coverslips) / 1,163 (Syn); +UK5099: 3 (Cultures) / 9 (Coverslips) / 230 (Syn) | Ctrl: 34.395 ± 0.977; +UK5099: 3.247 ± 0.61 | Two-Sample t-test |
| Figure 4I | 3-9 (culture), Ctrl: 9 (Cultures) / 36 (Coverslips) / 1,163 (Syn); +UK5099: 3 (Cultures) / 9 (Coverslips) / 230 (Syn) | Ctrl: 3.241 ± 0.094; +UK5099: 0.487 ± 0.068 | Two-Sample t-test |
| **Figure S1** | **Sample number** | **Mean ± SEM** | **Statistical test** |
| Figure S1A | 3 (cultures) | Ctrl: 1.0 ± 0.0; MPC1 KD: 0.08 ± 0.04 | Mann-Whitney U Test |
| **Figure S2** | **Sample number** | **Mean ± SEM** | **Statistical test** |
| Figure S2A | 3 (culture), Ctrl: 9 (wells), MPC1 KD:9 (wells) | Ctrl: 1 ± 0.0; MPC1 KD: 0.43 ± 0.06 | Mann-Whitney U Test |
| Figure S2B, C | 3 (culture), 6 (wells) | Ctrl: 1 ± 0.0 Sirt3 KD: 0.26 ± 0.09 | Mann-Whitney U Test |
| Figure S2D | 4 (culture), Ctrl: 7 (coverslips), 26 (FOV), 318 (cells); MPC1 KD : 5 (coverslips), 18 (FOV), 272 (cells), Sirt3 KD: 7 (coverslips), 25 (FOV), 388 (cells), #12-15 cells/ FOV | Ctrl: 1. ± 0.02, MPC1 KD: 1.0 ± 0.03, Sirt3 KD: 0.95 ± 0.01 | Kruskal-Wallis Test |
| **Figure S2** | **Sample number** | **Mean ± SEM** | **Statistical test** |
| Figure S3A, B | 3 (culture), Ctrl: 9 (coverslips/neurons), UK5099: 9 (coverslips/neurons) | Ctrl: 0.14 ± 0.02; UK5099: 0.59 ± 0.11 | Paired Wilcoxon Test |
| Figure S3C, D | 3 (culture), Ctrl: 17 (coverslips/neurons), Zaprinast: 17 (coverslips/neurons) | Ctrl: 0.13 ± 0.02; Zaprinast: 0.7 ± 0.07 | Paired Wilcoxon Test |
| Figure S3E, F | 4-13 (culture), Ctrl: 46 (coverslips), 81 (neurons) ; MPC1 KD: 35 (coverslips), 71 (neurons); MPC1 KD + WT: 24 (coverslips), 25 (neurons) | Ctrl: 0.14 ± 0.01; MPC1 KD: 0.39 ± 0.03; MPC1 KD + WT: 0.12 ± 0.02 | Kruskal-Wallis Test |

FOV: Field of view
Syn: synapses

Table S3
